# An experimental approach in analyzing the cell cycle dynamics of food-entrapping cells of sponges

**DOI:** 10.1101/2023.08.23.554503

**Authors:** Nikolai P. Melnikov, Andrey I. Lavrov

## Abstract

Sponges (phylum Porifera) exhibit surprisingly complex tissue dynamics, maintaining constant cell turnover and migration, rearranging internal structures, and regenerating after severe injuries. Such tissue plasticity relies on the activity of proliferating cells represented primarily by the food-entrapping cells, choanocytes. While there is plenty of studies regarding the dynamics of regeneration and tissue rearrangement in sponges, cell cycle kinetics of choanocytes in intact tissues remains a controversial issue.

This study is devoted to the comparative description of choanocyte cell cycle dynamics in intact tissues of two sponges, *Halisarca dujardinii* (class Demospongiae) and *Leucosolenia corallorrhiza* (class Calcarea). We have identified populations of proliferating cells and synchronized them in the S-phase to estimate the growth fraction of cycling cells. Using continuous exposure to labeled thymidine analog EdU, we calculated choanocyte cell cycle duration and the length of the S-phase. We also applied double labeling with EdU and antibodies against phosphorylated histone 3 to estimate the lengths of choanocyte M- and G2-phases. Finally, flow cytometry-based quantitative analysis of DNA content provided us with the lengths of G2- and G1-phases.

We found that tissue growth and renewal in studied sponges are generally maintained by a relatively large population of slowly cycling choanocytes with a total cell cycle duration of 40 hours in *H. dujardinii* and 60 hours in *L. corallorrhiza*. In both species, choanocytes are characterized by an extremely short M-phase and heterogeneity in the duration of the G2-phase.

## Introduction

Sponges (phylum Porifera) are an early-branching group of multicellular animals represented by aquatic sessile filter-feeding organisms of unique anatomy and physiology [1]. They lack tissues and organs typical of other animals, relying on constant pumping of surrounding water through a complex system of canals and chambers called the aquiferous system [2,3]. The central cell type in the aquiferous system are choanocytes, flagellated collar cells responsible for food uptake [3,4]. The aquiferous system is surrounded by the mesohyl, a combination of extracellular matrix, mineral skeleton, and scattered wandering cells of different origin and function.

Being sessile suspension feeders, sponges are in constant need to adapt to the ever-changing environment. Their tissues are highly plastic, constantly renewing under steady-state conditions and quickly regenerating after severe injuries [5–8]. Sponges are also able to rearrange their tissues according to the needs of the organism, consequently changing in shape, cell composition, and physiology [9,10]. Such a “chronic morphogenesis” [9] relies heavily on cellular plasticity manifesting in a form of transdifferentiation, i.e. the ability of some sponge cells to change their morphology and functions [2,4,11,12].

To maintain tissue homeostasis, continuous (trans)differentiation needs a source of new cells, namely a population of proliferating adult stem cells [13]. In sponges, certain amoeboid cells of mesohyl were traditionally thought to be pluripotent stem cells [4,14–16]. However, recent studies have shown choanocytes to play a leading role in regeneration [11,17–19] as well as in regular cell turnover [5,6,20,21]. If so, the dynamics of choanocyte proliferation should determine the general rate of cell turnover and regeneration speed. Most studies, however, are limited to the analysis of the proportion of proliferating cells rather than the length of the cell cycle.

Yet, the estimation of cell cycle length in animals can be quite a challenge. While there are various techniques of cell cycle analysis ranging from direct observation to the usage of different cell cycle-related reporters [22–24], they are mostly applicable to study cell cultures. Speaking of cell cycle in intact (and non-model) animals, we are usually constrained to using indirect methods of analysis implementing certain mathematical models [25,26]. Such studies regarding choanocytes of sponges provided us with a broad range of cell cycle estimates varying from 5.4 h in *Halisarca caerulea* [5] to 20-40 h in *Hymeniacidon sinapium* [27] and more than 150 h in *Spongilla lacustris*, *Haliclona mollis* and *Aphrocallistes vastus* [20].

Keeping in mind that choanocyte cell cycle length can differ significantly between species, we addressed this issue to two sponges belonging to distinct poriferan lineages and showing different anatomical organization, leuconoid *Halisarca dujardinii* (Class Demospongiae) and asconoid *Leucosolenia corallorrhiza* (Class Calcarea) (Fig. 1). While there are several papers devoted to regular cell turnover and regeneration dynamics in these species [11,19,21], the kinetics of choanocyte cell cycle has long been an unsettled issue. The intention of this study is to characterize proliferating cell populations and choanocyte cell cycle dynamics in intact tissues of *H. dujardinii* and *L. corallorrhiza*. In this paper, we use: a) a combination of thymidine analog EdU and cytoplasmic dye CellTracker to describe the morphology of proliferating cells; b) DNA-polymerase inhibitor aphidicolin to synchronize proliferating cells and estimate the growth fraction in intact animals; c) antibodies to phosphorylated histone 3 to describe the pattern of choanocyte mitoses and calculate the length of M- and G2-phases; d) analysis of EdU incorporation dynamics to calculate the duration of the cell cycle; e) quantitative DNA staining and flow cytometry to determine the length of individual cell cycle phases. Our data not only complement the general knowledge of the morphogenetic processes in sponges but also provide a blueprint for studying the cell cycle in other animals.

**Fig. 1.**
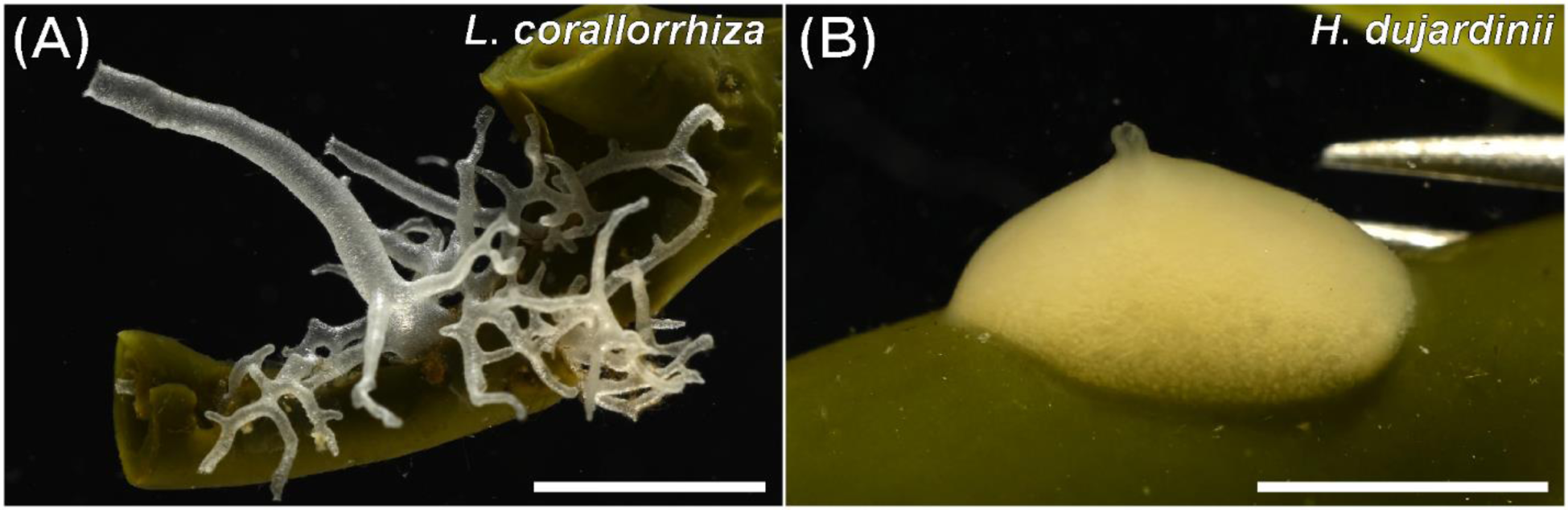
Representatives of studied sponges. A – intact *L. corallorrhiza* attached to a brown algae. B – intact *H. dujardinii* attached to a brown algae. Scale bars: A, B – 5 mm.

## Results

### Identifying proliferating cell types

We used the combination of labeled nucleotides EdU and cytoplasmic dye CellTracker to describe the morphology of proliferating cells in intact tissues of demosponge *Halisarca dujardinii* and calcareous sponge *Leucosolenia corallorrhiza*. Both species show a considerable number of cycling cells (Figs. 2B, 3B) with total labeling fraction (the proportion of all EdU-positive nuclei to the total cell number; mean ± SE) reaching 5.3±0.9% in *H. dujardinii* and 9.2±0.4% in *L. corallorrhiza*. The labeling fraction of choanocytes (the proportion of EdU-positive choanocytes to the number of choanocytes) was 10.5±1.8% and 13.4±1.0%, respectively (Table 1). As we reported earlier [21], the absolute majority (>90%) of labeled cells in both species are represented by choanocytes, flagellated food-entrapping cells of the aquiferous system. Minor populations of cycling cells, however, differ between species, probably reflecting the difference in the anatomical and histological structure of studied sponges.

**Fig. 2.**
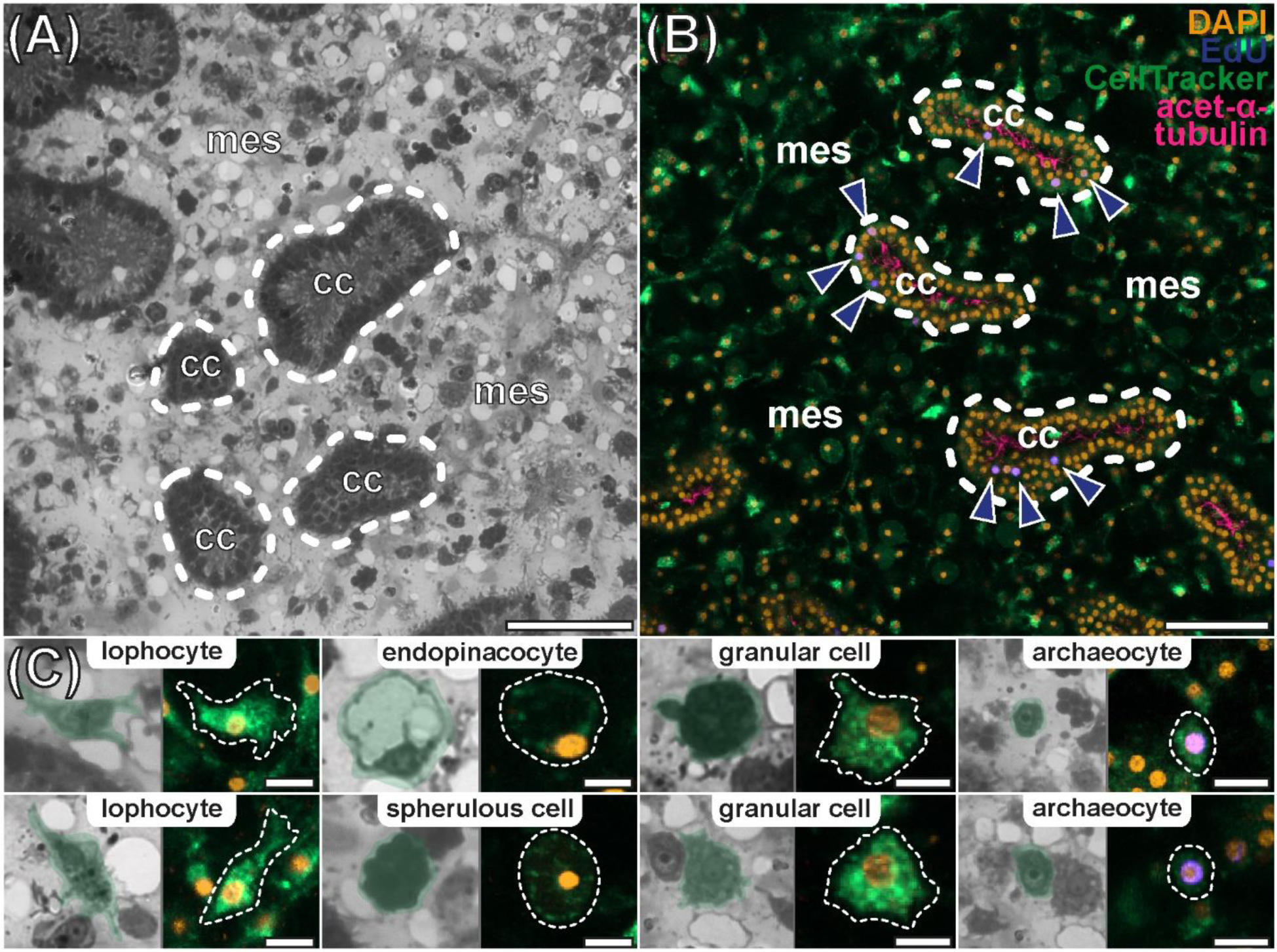
Cell proliferation in intact tissues of *H. dujardinii*. A – general view of the endosome, semi-thin section. B – proliferation in the endosome, CLSM. C – cell types studied by comparing semi-thin sections to cytoplasmic CellTracker staining. cc – choanocyte chambers, mes – mesohyl. Blue arrowheads mark EdU-labeled cells. Orange – DAPI, cell nuclei; blue – EdU, nuclei of DNA-synthesizing cells; green – CellTracker, cytoplasmic dye; magenta – acetylated α-tubulin, flagella. Scale bars: A, B – 50 µm; C – 10 µm.

**Fig. 3.**
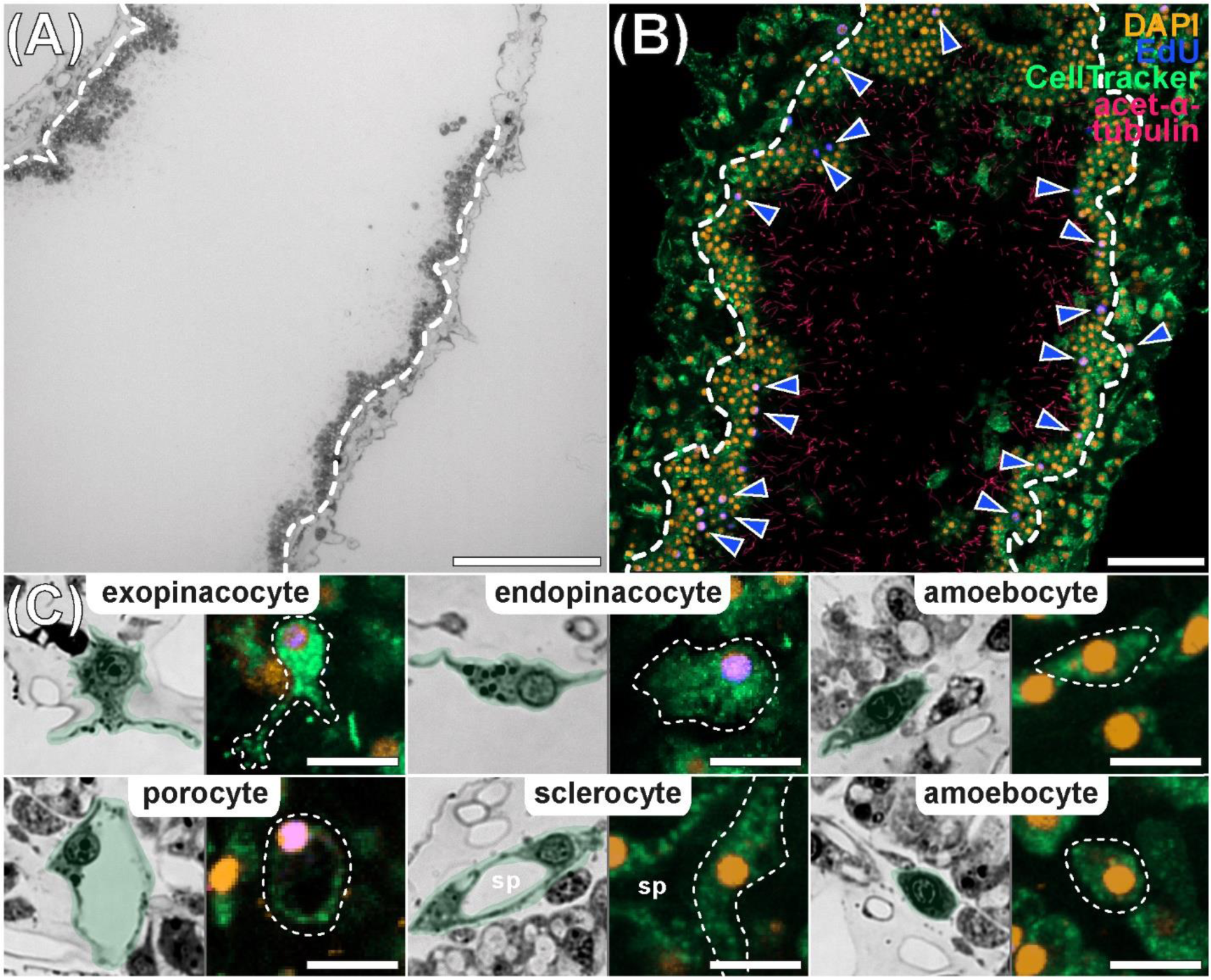
Cell proliferation in intact tissues of *L. corallorrhiza*. A – general view of the oscular tube, semi-thin longitudinal section. B – proliferation in the middle part of the oscular tube, CLSM. C – cell types studied by comparing semi-thin sections to cytoplasmic CellTracker staining. sp – volume previously occupied by a spicule. Blue arrowheads mark EdU-labeled cells. Orange – DAPI, cell nuclei; blue – EdU, nuclei of DNA-synthesizing cells; green – CellTracker, cytoplasmic dye; magenta – acetylated α-tubulin, flagella. Scale bars: A, B – 50 µm; C – 15 µm.

**Table 1.**
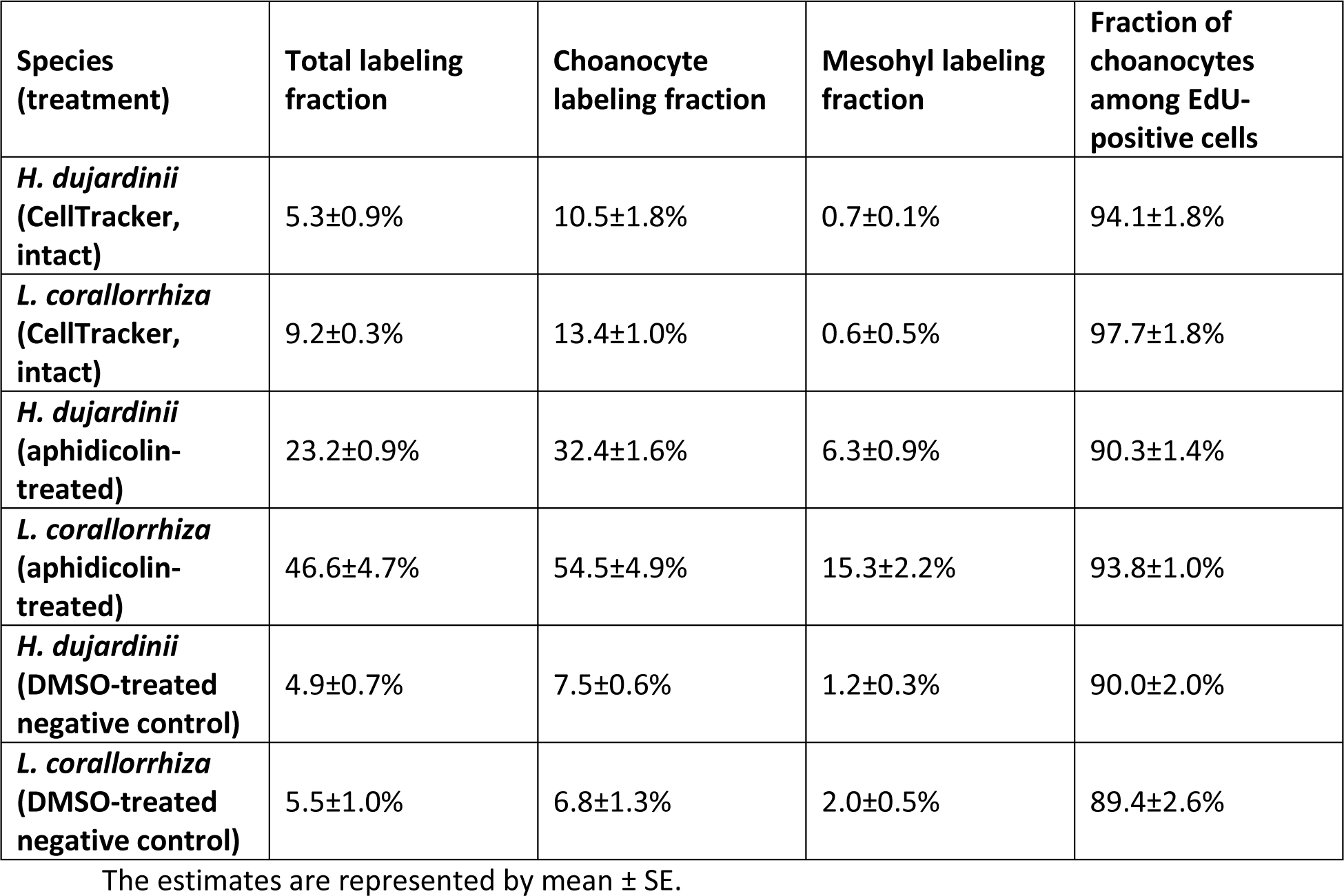
The calculated fractions of labeled cells in different sets of experiments, CLSM.

| Species (treatment) | Total labeling fraction | Choanocyte labeling fraction | Mesohyl labeling fraction | Fraction of choanocytes among EdU-positive cells |
| --- | --- | --- | --- | --- |
| *H. dujardinii* (CellTracker, intact) | 5.3±0.9% | 10.5±1.8% | 0.7±0.1% | 94.1±1.8% |
| *L. corallorrhiza* (CellTracker, intact) | 9.2±0.3% | 13.4±1.0% | 0.6±0.5% | 97.7±1.8% |
| *H. dujardinii* (aphidicolin-treated) | 23.2±0.9% | 32.4±1.6% | 6.3±0.9% | 90.3±1.4% |
| *L. corallorrhiza* (aphidicolin-treated) | 46.6±4.7% | 54.5±4.9% | 15.3±2.2% | 93.8±1.0% |
| *H. dujardinii* (DMSO-treated negative control) | 4.9±0.7% | 7.5±0.6% | 1.2±0.3% | 90.0±2.0% |
| *L. corallorrhiza* (DMSO-treated negative control) | 5.5±1.0% | 6.8±1.3% | 2.0±0.5% | 89.4±2.6% |

The estimates are represented by mean ± SE.

*Halisarca dujardinii* is a thick globular sponge with a leuconoid aquiferous system. The inner area of its body – the endosome – contains numerous choanocyte chambers (Fig. 2A). Choanocyte chambers are connected via canals and surrounded by an extensive, well-developed mesohyl containing several cell types (Fig. 2A). Since the morphology of these cells was described earlier [28], we tried to identify them in confocal Z-stacks by comparing CellTracker staining to semi-thin histological sections (Fig. 2C). We were able to distinguish elongated collagen-secreting lophocytes, cylindrical endopinacocytes forming canals, spherulous cells with small nuclei and large inclusions, and a heterogeneous population of amoeboid cells. Amoeboid cells were subdivided into two groups, one being represented by large (9-14 µm in diameter) cells of irregular shape, and the other – by small (6-9 µm in diameter) oval or elongated cells with a thin rim of cytoplasm surrounding the nucleus (Fig. 2C). We suppose that the larger cells are granular cells (amoebocytes with inclusions), while the smaller cells are archaeocytes (presumptive pluripotent mesohyl cells). The latter were the only EdU-labeled cells outside the choanoderm after 12 h of EdU treatment (Fig. 2C).

*Leucosolenia corallorrhiza* is an asconoid sponge composed of a plexus of thin-walled tubes called cormus. Cormus gives rise to multiple oscular tubes used to expel filtered water. The interior of oscular and cormus tubes is lined with a continuous layer of choanocytes, while T-shaped exopinacocytes make up the outer layer of cells (Fig. 3A, B). Both cell layers are connected with ostia formed by porocytes (Fig. 3C). Mesohyl is located between the exopinacoderm and choanoderm. The mesohyl contains few cells, mostly spicule-secreting sclerocytes and cells with amoeboid morphology [29] (Fig. 3C). Nuclei of porocytes, exopinacocytes, sclerocytes and amoebocytes are scattered randomly in the mesohyl, making the CellTracker staining essential to distinguish between EdU-labeled cell types (Fig. 3B). After 6 h of EdU treatment, the population of EdU-labeled cells outside the choanoderm consisted mostly of exopinacocytes and porocytes (Fig. 3C). The rest of labeled cells exhibited irregular shape and amoeboid-like morphology; these, however, could – at least partly – also be cell bodies of T-shaped exopinacocytes since they often form pseudopodia-like protrusions [29] (Fig. 3C). The last type of EdU-positive cells were flat endopinacocytes which replace choanocytes in the oscular rim (the most distal part of the oscular tube) (Fig. 3C).

### Estimating the growth fraction

A short EdU treatment provides nothing more than a snapshot of DNA-synthesizing (i.e. S-phase) cells; G1- and most G2-cells usually escape this kind of analysis. To reveal these “hidden” cells, we used an aphidicolin-mediated S-phase block. Since aphidicolin inhibits DNA synthesis, continuous treatment with the drug results in the accumulation of proliferating cells just at the beginning of the S-phase. Given the treatment time exceeds the duration of the cell cycle, it is possible to accumulate all cycling cells (the growth fraction) at the beginning of the S-phase and label them with EdU.

As expected, 4.5 days of aphidicolin treatment drastically increased the number of labeled cells (Fig. 4A). The total labeling fraction (the proportion of all EdU-positive nuclei to the total cell number; mean ± SE) reached 23.2±0.9% in *H. dujardinii* and 46.6±4.7% in *L. corallorrhiza* (Fig. 4B, Table 1). The difference between the total labeling fraction in DMSO-treated sponges (negative controls) and aphidicolin-treated sponges was significant with Wald and log-likelihood p-values being less than 0.001. The labeling fraction of choanocytes (the proportion of EdU-positive choanocytes to the number of choanocytes) was 32.4±1.6% in *H. dujardinii* and 54.5±4.9% *L. corallorrhiza* (Fig. 4B, Table 1), while mesohyl contained 6.3±0.9% and 15.3±2.2% labeled cells, respectively (Fig. 4B, Table 1). Wald and likelihood tests reported p-values less than 0.001 for the effect of aphidicolin treatment on the choanocyte and mesohyl labeling fractions. The fraction of choanocytes among all EdU-positive cells remained the same as in intact tissues, reaching 90.3±1.4% in *H. dujardinii* and 93.8±1.0% in *L. corallorrhiza* (Table 1).

**Fig. 4.**
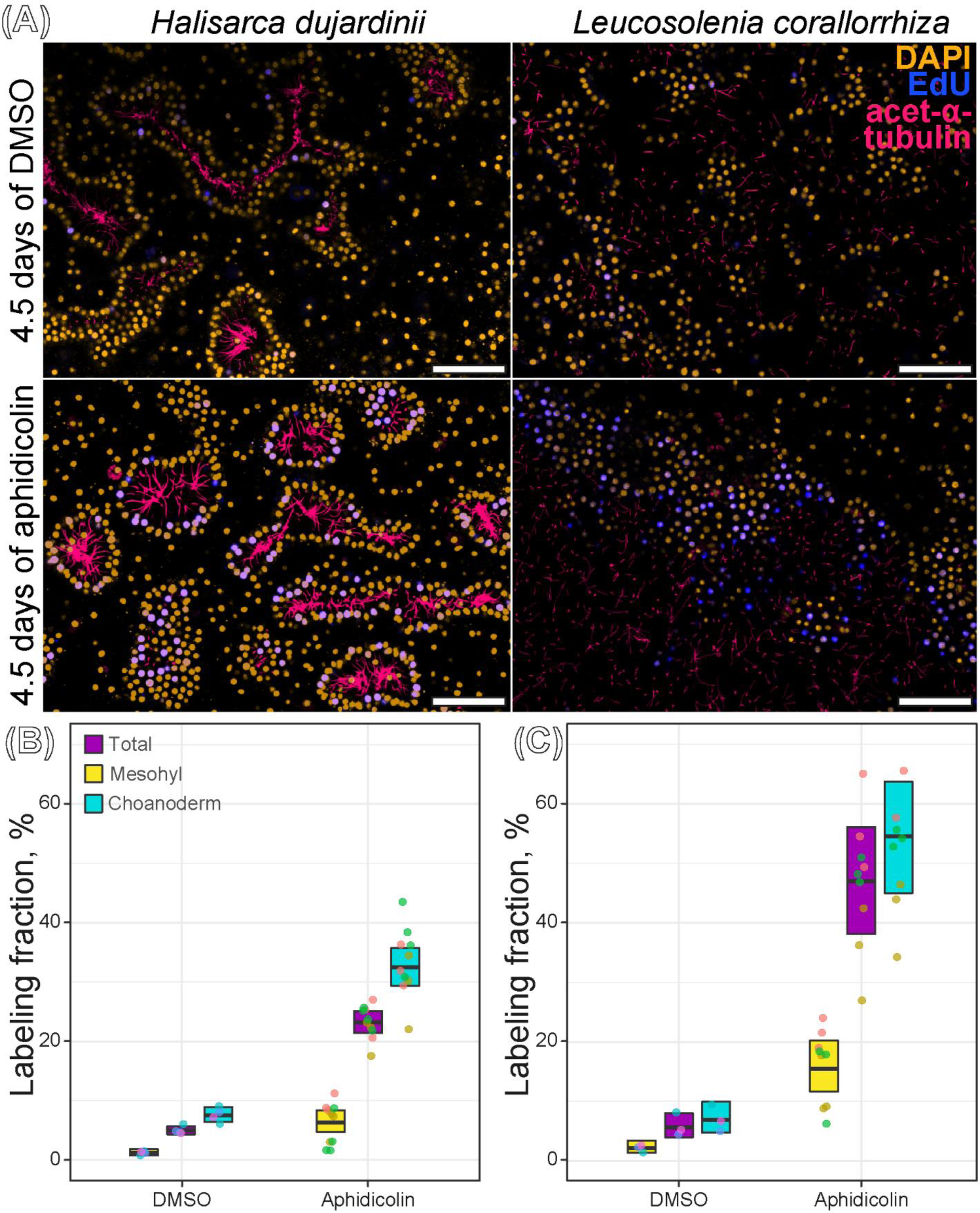
Aphidicolin-mediated S-phase synchronization of choanocytes, CLSM. A – confocal images of aphidicolin- and DMSO-treated sponge tissues. Orange – DAPI, cell nuclei; blue – EdU, nuclei of DNA-synthesizing cells; magenta – acetylated α-tubulin, flagella. B – quantitative analysis of synchronization in *H. dujardinii*. C – quantitative analysis of synchronization in *L. corallorrhiza*. Total labeling fraction is the proportion of all EdU-positive cells to the total number of cells; choanocyte labeling fraction is the proportion of EdU-positive choanocytes to the total number of choanocytes; mesohyl labeling fraction is the proportion of EdU-positive mesohyl cells to the total number of mesohyl cells. Data are shown with individual values (dots), mean values (thick horizontal lines), and 95% confidence intervals (boxes). Dots of the same color represent values obtained from different Z-stacks of the same individual. Scale bars: A – 50 µm.

Notably, sponges treated with DMSO instead of aphidicolin (negative controls) showed fewer labeled cells when compared to the previous experiments (see “Identifying proliferating cell types” section). DMSO-treated *L. corallorrhiza* had the total labeling fraction of 5.5±1.0% (compared to 9.2±0.3% in intact sponges) and choanocyte labeling fraction of 6.8±1.3% (compared to 13.4±1.0%) (Table 1). In *H. dujardinii* the total labeling fraction remained at the same level of 4.9±0.7% (compared to 5.3±0.9% in intact sponges), but choanocyte labeling fraction slightly decreased to 7.5±0.6% (compared to 10.5±1.8%) (Table 1). Sponges exposed to EdU before the end of aphidicolin treatment (positive controls) still contained labeled cells; these cells were, however, extremely faint and, in most cases, very few.

Thus, we consider the growth fraction of choanocytes (the fraction of all cycling choanocytes) to be 32.4±1.6% in *H. dujardinii* and 54.5±4.9% in *L. corallorrhiza*.

### G2/M-phase analysis

We used antibodies against Ser10-phosphorylated histone 3 to mark cells in the late G2- and M-phases and analyze the orientation of division planes. Studied sponges had few pH3-positive cells accounting for 0.56±0.13% (mean ± SE) of the total cell number in *H. dujardinii* and 0.35±0.05% in *L. corallorrhiza*. Choanocytes comprised 84.6±2.7% of pH3-positive cells in *H. dujardinii* and 82.7±1.7% in *L. corallorrhiza*. The mitotic index in the choanoderm was 0.64±0.15% in *H. dujardinii* and 0.4±0.07% in *L. corallorrhiza*. Judging by the orientation of mitotic plates and fission spindles, all registered instances of choanocyte mitoses were symmetric with both daughter cells remaining in the choanoderm after the division (Fig. 5A-C).

**Fig. 5.**
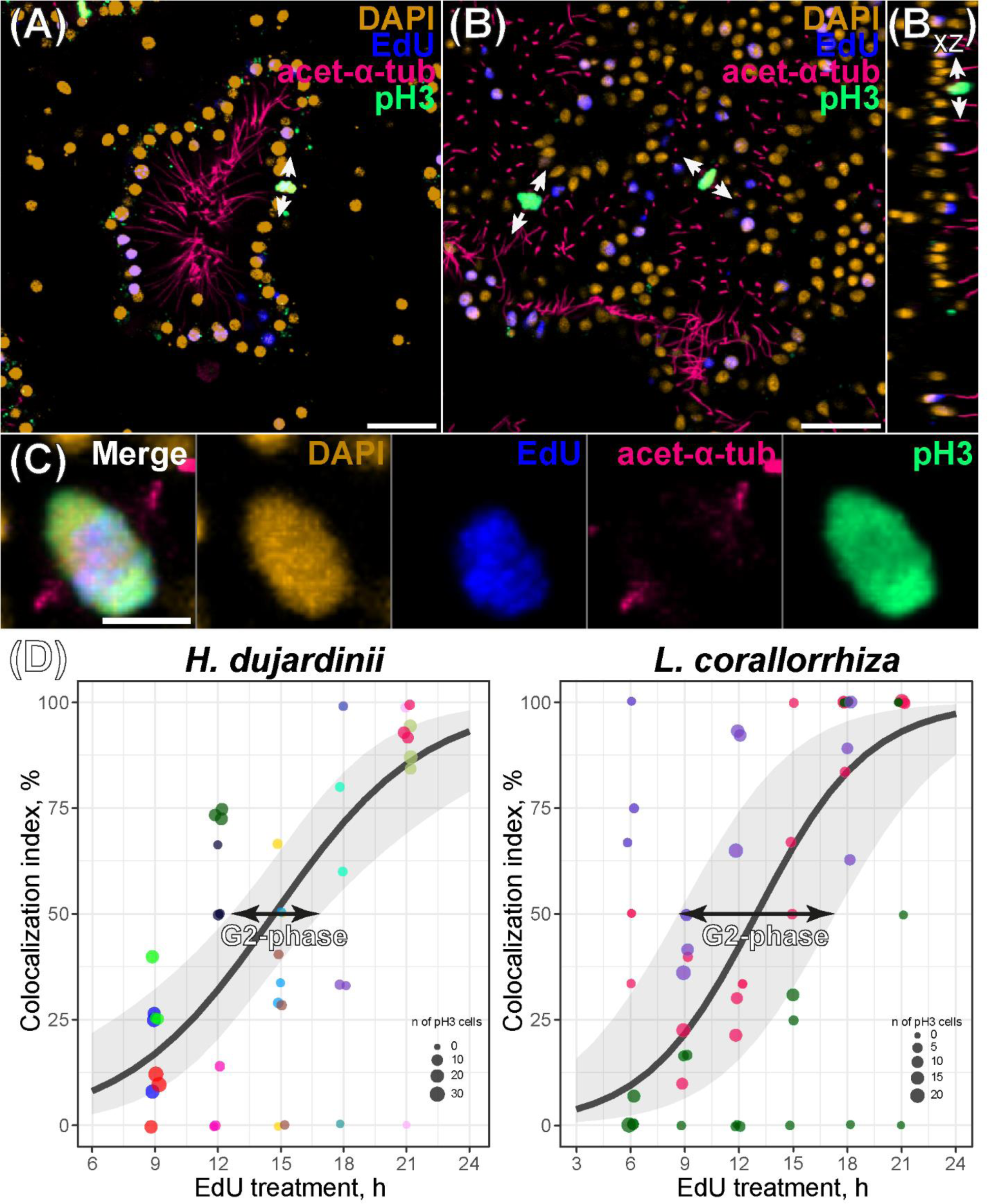
G2/M-phase analysis of choanocytes. A – mitosis in a choanocyte chamber of *H. dujardinii*, CLSM; arrows indicate the orientation of the mitotic spindle. B – mitoses in choanoderm of *L. corallorrhiza*, CLSM; arrows indicate the orientation of the mitotic spindle. C – an example of EdU-labeled pH3-positive choanocyte in *L. corallorrhiza*, CLSM. Orange – DAPI, cell nuclei; blue – EdU, nuclei of DNA-synthesizing cells; magenta – acetylated α-tubulin, flagella; green – phosphorylated histone 3, M- or late G2-phase cells. D – quantitative analysis of the accumulation of EdU-labeled pH3-positive choanocytes in *H. dujardinii* and *L. corallorrhiza* during continuous EdU treatment. Colocalization index is the fraction of labeled mitotic cells (i.e., the proportion of EdU-positive pH3-positive choanocytes to the total number of pH3-positive choanocytes). Data are shown with individual values (dots), predicted mean values (black line), and 95% confidence interval of prediction (transparent grey band). Dots of the same color represent values obtained from different Z-stacks of the same individual. Scale bars: A, B – 30 µm; C - 5 µm.

We also tried to track the transition of choanocytes through the G2-phase by the accumulation of EdU-positive mitotic cells during the continuous EdU treatment. Labeled mitoses were already present at 9 h (*H. dujardinii*) and 6 h (*L. corallorrhiza*) after the beginning of the EdU treatment (Fig. 5D). They were, however, few, and accounted for 6.2±3.8% and 7.0±4.9% of all registered mitoses, respectively (Fig. 5). The fraction of labeled mitoses eventually increased but seldom reached 100%. Nonetheless, regression analysis revealed strong effect of the EdU treatment duration on the fraction of labeled mitoses with p-values of log-likelihood and Wald tests being less than 0.001 in both species. Regression-derived estimates of the fraction of labeled mitoses were 85.3±6.1% (*H. dujardinii*) and 90.7±6.3% (*L. corallorrhiza*) at the 21^st^ h of the experiment. According to the 95% confidence intervals of the prediction, the average duration of the G2-phase (time point with colocalization index reaching 50%) was between 8 and 13 hours in *H. dujardinii* and 9 and 17 hours in *L. corallorrhiza* (Fig. 5D). The average marginal effect of the EdU treatment was estimated as 5.0±0.7% of mitoses becoming labeled with the increase of EdU treatment duration by one hour in *H. dujardinii* and 4.9±0.3% in *L. corallorrhiza*.

### Cell cycle dynamics

The estimation of choanocyte cell cycle dynamics was performed by analyzing the kinetics of EdU incorporation. We treated sponges with EdU for **T_E1_**=2 h and **T_E2_**=6 h and observed an increase in the labeling fraction of choanocytes (Fig. 6). After 2 h of treatment, the mean choanocyte labeling fraction ± SE (**F_E1_**) was 11.4±0.8% in *H. dujardinii* and 13.7±0.9% in *L. corallorrhiza*; 6 h of treatment resulted in the labeling fraction **F_E2_** of 14.6±1.0% in *H. dujardinii* and 17.4±1.2% in *L. corallorrhiza* (Fig. 6). Wald and log-likelihood tests showed the support for the increase in the labeling fractions of choanocytes with the respective p-values being 0.0151 and 0.04397 in case of *H. dujardinii* and both less than 0.0001 in *L. corallorhiza*. The estimated average marginal effect of the EdU treatment (meaning the fraction of choanocytes becoming labeled each hour of incubation) was 0.8±0.3% in *H. dujardinii* and 0.9±0.1% in *L. corallorrhiza*.

**Fig. 6.**
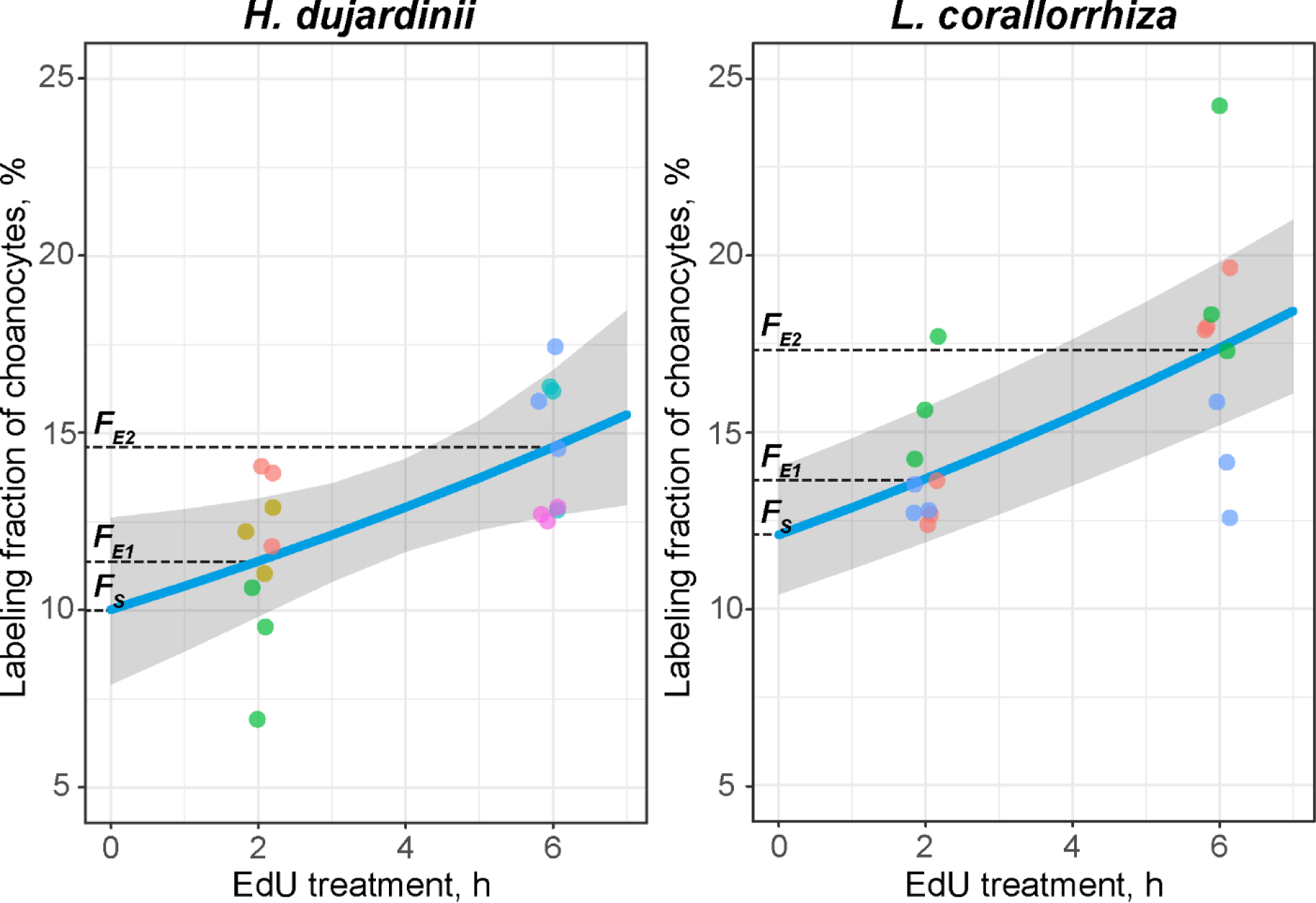
The accumulation of EdU-labeled choanocytes during the short-term continuous EdU treatment. Data are shown with individual values (dots), predicted mean values (blue line), and 95% confidence interval of prediction (transparent grey band). Dots of the same color represent values obtained from different Z-stacks of the same individual. F_E1_ is the labeling fraction after 2 h of treatment; F_E2_ is the labeling fraction after 6 h of treatment; F_S_ is the intercept of the logistic regression (the fraction of choanocytes in the S-phase).

Then, we applied Sanders’ mathematical model to calculate the duration of the cell cycle **T_C_** using the following formula with growth fraction (**GF**) being equal to the labeling fraction of choanocytes after the aphidicolin-mediated synchronization (see “Estimating the growth fraction” section): 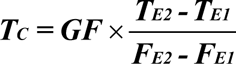. The total duration of the cell cycle **T_C_** was 40.3±16.9 h in *H. dujardinii* and 58.7±24.7 h in *L. corallorrhiza* (the estimates are represented as mean ± SE) (Table 2).

**Table 2.** The calculated parameters of choanocyte cell cycle in *H. dujardinii* and *L. corallorrhiza*.

| Species | Total cell cycle length $T_E$ | S-phase duration $T_S$ | M-phase duration $T_E$ | G2-phase duration $T_{G2}$ (CLSM/flow cytometry) | G1-phase duration $T_{G1}$ |
| --- | --- | --- | --- | --- | --- |
| *H. dujardinii* | 40.3±16.9 h | 12.5±5.5 h | 0.7±0.3 h | 8-13 / ~15 h | ~13 h |
| *L. corallorrhiza* | 58.7±24.7 h | 13.1±5.7 h | 0.4±0.2 h | 9-17 / ~18 h | ~28 h |

The estimates are represented by mean ± SE.

The intercept of logistic regression (the fraction of choanocytes in S-phase at any given moment, **F_S_**) was 10.0±2.3% in *H. dujardinii* and 12.1±1.8% in *L. corallorrhiza* (Fig. 6). We supposed that the **F_S_** reflected the relative duration of the S-phase (as usually occurs in asynchronous cycling cell populations; [22]) and used the following formula to calculate the length of the S-phase 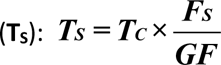. The duration of S-phase was 12.5±5.5 h in *H. dujardinii* and 13.1±5.7 h in *L. corallorrhiza* (Table 2). The length of the M-phase (**T_M_**) was calculated in a similar fashion using the mitotic fraction in choanoderm (**F_M_**) provided earlier (see the “G2/M-phase analysis” section): 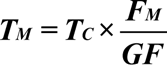. The estimated length of the M-phase was 0.7±0.3 h in *H. dujardinii* and 0.4±0.2 h in *L. corallorrhiza* (Table 2).

### Flow cytometry: cell cycle distribution

We used propidium iodide (PI) staining to analyze DNA content and assess cell cycle distribution in single-cell suspensions of studied sponges, providing an alternative way to estimate the length of choanocyte G2-phase apart from colocalization studies. In general, both species show similar DNA content curves with most of the cells (47.9±11.8% in *H. dujardinii* and 33.5±7.2% in *L. corallorrhiza*; mean ± SD) belonging to a diploid G0/G1 cell population (Fig. 7). Suspensions also contained a considerable number of tetraploid G2/M cells reaching 14.7±2.1% in *H. dujardinii* and 15.3±1.4% in *L. corallorrhiza*. Cells in the S-phase accounted for 12.0±2.1% and 11.2±1.6% cells, respectively (Fig. 7). Considering that more than 90% of proliferating cells in intact tissues were choanocytes (see “Identifying proliferating cell types” and “Estimating the growth fraction” sections), the absolute majority of S- and G2/M-cells should be choanocytes, too. We used this assumption to qualitatively estimate the lengths of choanocyte G1- and G2-phases.

**Fig. 7.**
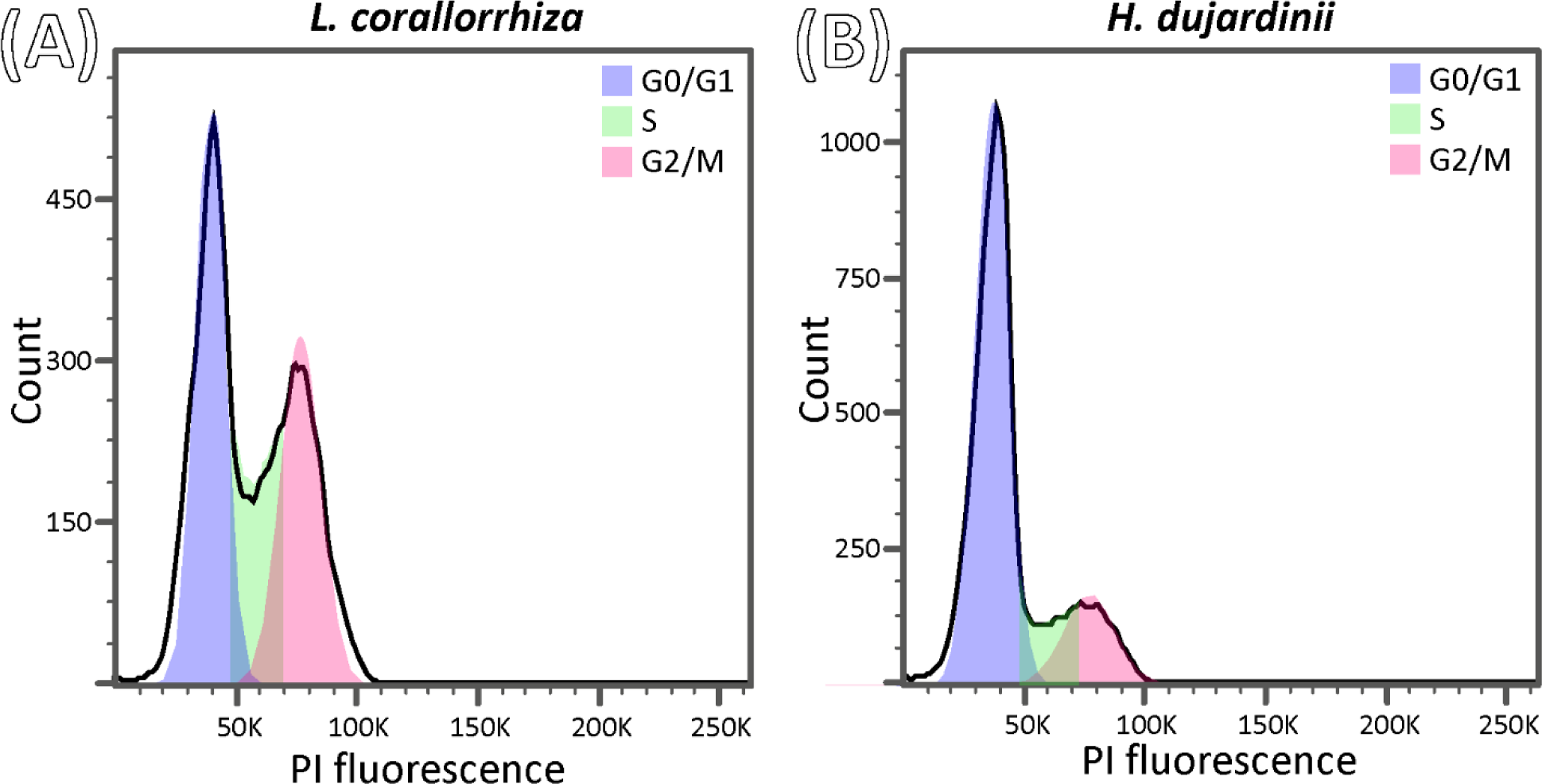
Quantitative DNA staining of sponge cell suspensions, flow cytometry. A – single-parameter histogram of PI fluorescence (i.e. DNA content) in cell suspensions of *L. corallorrhiza*. B – single-parameter histogram of PI fluorescence (i.e. DNA content) in cell suspensions of *H. dujardinii*. G0/G1 (blue) are diploid cells, G2/M (pink) are tetraploid cells, S (green) are DNA-synthesizing cells.

The mean proportion of G2- and M-cells to S-cells (G2/M : S ratio) was 1.2 in *H. dujardinii* and 1.4 in *L. corallorrhiza*, indicating combined G2- and M-phases to be slightly longer than the S-phase. Since we found pH3-positive cells to be few in intact tissues (see “G2/M-phase analysis” section), the duration of the M-phase is quite negligible, and the majority of G2/M-cells are actually cells in the G2-phase. Given the S-phase duration was estimated earlier to be 12.5 h (*H. dujardinii*) and 13.1 h (*L. corallorrhiza*), the corresponding length of the G2-phase of choanocytes (**T_G2_**) should be about 15 h in *H. dujardinii* and 18 h in *L. corallorrhiza* (Table 2). Knowing the total length of the cell cycle (**T_C_**) and the lengths of S-, G2- and M-phases (**T_S_**, **T_G2_**, and **T_M_**, respectively), we were able estimate the length of G1-phase: ***T_G1_ = T_C_ - T_S_ - T_G2_ - T_M_***. Choanocyte G1-phase was 13 h in *H. dujardinii* and 28 h in *L. corallorrhiza* (Table 2).

## Discussion

### Sponges possess several types of proliferating cells

Sponges are sessile aquatic animals well-known for their strikingly dynamic tissues. One of the major components of animal tissue dynamics is cell proliferation, usually maintained by somatic stem cells [13,30]. There has long been a debate about which cell type acts as the primary stem cells in sponges. While certain amoeboid cells of mesohyl were traditionally regarded as the main type of stem cells in sponges [2,4,14,31], recent studies [6,20,21] showed that the majority of proliferating cells in intact tissues is actually comprised of food-entrapping cells of the aquiferous system, choanocytes. Here we also show that choanocytes are the main proliferating cell type both in intact sponges and after the S-phase synchronization.

Notably, choanocytes are actively cycling in sponges belonging to different Porifera lineages – namely, in *Halisarca dujardinii* (class Demospongiae) and *Leucosolenia corallorrhiza* (class Calcarea). Choanocytes were also reported to proliferate in other demosponges [5,6], calcareous sponges [20], hexactinnelids [20], and homoscleromorphs [17], thus representing a common type of cycling cells in sponges.

Still, choanocytes are not the only cycling cells in sponges. As expected, we found archaeocytes (amoeboid mesohyl cells) to incorporate labeled nucleotides in demosponge *H. dujardinii*; in calcareous sponge *L. corallorrhiza*, however, proliferating amoebocytes are few (if any), and cycling cells outside the choanoderm are represented by epithelial-like exopinacocytes, endopinacocytes, and ostia-forming porocytes. In general, the presence of cycling mesohyl cells seems to be a common feature of demosponges and hexactinnelids, while the proliferation of epithelial-like cycling cells is characteristic of Calcarea + Homoscleromorpha group (e.g., exopinacocytes were shown to incorporate EdU in homoscleromorph *Oscarella lobularis*) [17].

### S-phase synchronization reveals numerous cycling cells in choanoderm

In studied species, cycling cells outside the choanoderm are extremely few, and so far there is no evidence that they proliferate much faster than choanocytes. If so, the general tissue dynamics in intact sponges is determined by the rate of choanocyte proliferation. We suppose that the constant production of new choanocytes mainly helps to maintain the homeostasis of filtering structures. Here we used aphidicolin-mediated S-phase synchronization to show that the renewal of choanoderm in studied sponges depends on a relatively large (30-50% of the total cell number) population of cycling cells. The fraction of EdU-labeled choanocytes after the synchronization was considered as the growth fraction of choanoderm and used in the subsequent cell cycle analysis. Apparently, not all choanocytes are equally proliferative; there is a considerable number of non-cycling choanocytes. Although there could be some quiescent choanocytes not registered by our technique, we suppose that registered presumably non-cycling choanocytes might be “differentiated” cells residing in G0-phase; there is already some data indicating the presence of different subpopulations among choanocytes [32].

To the best of our knowledge, this is the first attempt to experimentally synchronize cycling cells in sponges. Generally, aphidicolin proved to be a promising tool to reversibly block proliferation without severely affecting the viability of animals. There are, however, certain matters of concern. Firstly, we found that it was insufficient to expose animals to EdU immediately after the aphidicolin treatment; it seems that sponges need some time to recover after the S-phase block. Secondly, DMSO-treated sponges (i.e., negative controls) showed fewer cycling cells when compared to intact sponges, probably reflecting the suboptimal maintenance conditions during such a prolonged experiment. Suboptimal conditions could well affect aphidicolin-treated sponges, too; in such case, the real number of cycling cells (meaning the growth fraction) in studied sponges could be even bigger than we observed. Thirdly, the presence of rare EdU-labeled cells in positive controls (sponges that were exposed to EdU before the end of aphidicolin treatment) indicates that aphidicolin – at least in a concentration of 1 µg/ml – is not always sufficient to entirely block the proliferation. Of course, this could also lead to the underestimation of the growth fraction. However, the intensity of EdU staining in these “aphidicolin-resistant” cells was usually quite low, hinting that aphidicolin still affected these cells. We suppose that occasionally aphidicolin just severely slowed the progression through the S-phase rather than entirely blocking it.

### Choanocytes possess a relatively long cell cycle

It is not only the number of proliferating choanocytes that affects the rate of cell turnover; the other matter worth paying attention to is the temporal characteristics of the cell cycle. There were several attempts to analyze the cell cycle dynamics of choanocytes resulting in a total cell cycle duration of 5.4 h in *Halisarca caerulea* (Demospongiae), 20-40 h in *Hymeniacidon synapium* (Demospongiae), 30 h in *Sycon coactum* (Calcarea), and nearly 150 h in *Spongilla lacustris* (Demospongiae), *Haliclona mollis* (Demospongiae) and *Aphrocallistes vastus* (Hexactinellida) [5,20,27]. Most of these studies were based on the Nowakowsky mathematical model analyzing the accumulation of labeled cells during the continuous exposure to labeled nucleotides [25]. Basically, each cell entering the S-phase become labeled, resulting in the increase in the labeling fraction. At a certain time point, all cells capable of proliferation pass through the S-phase, making the labeling fraction come to a plateau. The plateau serves as the estimate of the growth fraction, while the dynamics of the accumulation of labeled cells provide temporal characteristics of the cell cycle.

However, the increase in the labeling fraction can also occur due to the division of cells. Both daughter cells inherit the nucleotides incorporated during the S-phase, so each labeled cell produces two labeled cells after finishing the M-phase. Using antibodies against Ser10-phosphorylated histone 3, we showed that in both species choanocytes divide symmetrically with each daughter cell remaining in the choanoderm. Although there is some evidence that choanocytes eventually leave choanoderm and transdifferentiate into mesohyl cells [21,33], recently divided choanocytes are still able to account for the accumulation of labeled cells and bias the analysis. This fact should be kept in mind when using continuous treatment with labeled nucleotides.

In this paper, we introduce an alternative way to determine the cell cycle parameters of choanocytes. Initially, we checked the minimal length of choanocyte G2-phase by analyzing the incorporation of labeled nucleotides (EdU) into pH3-positive (mitotic) cells. To our surprise, few EdU-labeled pH3-positive choanocytes were already present after only 6 h of EdU treatment in *L. corallorrhiza* and 9 h – in *H. dujardinii*. These cells managed to finish S-phase and pass through the G2-phase during the exposure, marking the minimal length of the G2-phase to be around 6 h.

Then, we observed an increase in a fraction of labeled choanocytes between 2 and 6 hours of EdU treatment; 6 hours of exposure were chosen as the second time point in order to achieve the noticeable increase without cell division significantly affecting the accumulation. We supposed that the observed increase was mostly a result of choanocytes entering the S-phase and used Sanders’ mathematical model [34] to calculate the length of the cell cycle. Since not all choanocytes are cycling, we adjusted the formula to account for the growth fraction estimated in S-phase synchronization experiments.

The total length of choanocyte cell cycle was about 40 h in *H. dujardinii* and 60 h in *L. corallorrhiza* (Table 2). The difference between the estimates was mostly a consequence of *L. corallorrhiza* having larger growth fraction, since the number of choanocytes entering the S-phase each hour was nearly the same in *H. dujardinii* and *L. corallorrhiza*. Basically, *H. dujardinii* maintains the same rate of cell renewal as *L. corallorrhiza* despite having fewer cycling cells.

To sum up, cell turnover in the aquiferous systems of *H. dujardinii* and *L. corallorrhiza* is maintained by relatively large populations of proliferating cells. In fact, having a considerable number of cycling cells is fairly common in metazoan epithelial tissues such as the gills and the mantle of mollusks, insect intestine, cnidarian epithelia, and the intestinal epithelium of mammals [30,35]. Speaking of cell cycle dynamics, it was previously reported that in demosponge *Halisarca caerulea* the whole cell cycle lasted for only 5.4 hours [5]; on the contrary, choanocytes of *Spongilla lacustris* (Demospongiae), *Haliclona mollis* (Demospongiae) and *Aphrocallistes vastus* (Hexactinellida) were assumed to have cell cycle as long as 150 hours [20]. Our results are more consistent with cell cycle lengths known for *Hymeniacidon synapium* (20-40 hours) and *Sycon coactum* (30 hours) [20,27]. We suppose that the growth fraction in *H. dujardinii* and *L. corallorrhiza* consists of relatively slowly cycling choanocytes. Quite a long cell cycle seems characteristic of proliferating cells in adult animals. For instance, the cell cycle of epithelial and interstitial cells in *Hydra* takes up to 72 and 30 hours, respectively [36,37]; gill cells of *Mytilus galloprovincialis* have the cell cycle of 24-30 hours [38]; various proliferative cells of adult mice and rats often have cell cycle far longer than 24 hours [40]; finally, planarian neoblasts also seem to have prolonged cell cycle [41].

### Cell cycle parameters of choanocytes

We also tried to calculate the lengths of individual cell cycle phases of choanocytes. Since we found no waves of proliferation occurring in the choanoderm of studied sponges, we consider choanocytes as an asynchronous population of cycling cells. Thus, the proportion of choanocytes in a certain cell cycle phase reflects its duration [22]. Since we found few pH3-positive (i.e., mitotic) choanocytes, we consider the M-phase of choanocytes to take up less than 2% of the total cell cycle duration and last no more than 1 hour (Table 2). We believe that the short duration of choanocyte M-phase helps to balance between the needs for steady proliferation and constant water pumping. We also derived the proportion of choanocytes in the S-phase at any given time moment by using binomial regression. The S-phase accounts for about 31% of the cell cycle in choanocytes of *H. dujardinii* and 22% in *L. corallorrhiza* and lasts for 12 and 13 hours, respectively (Table 2).

The G2/M-phase analysis provided us with conflicting results. We expected to observe fast and linear increase in the fraction of labeled mitoses during continuous treatment. When the exposure time is less than the duration of the G2-phase, EdU-labeled cells do not have enough time to pass the G2-phase and begin the mitosis; thus, no EdU-labeled pH3-positive (i.e., mitotic) cells are present (the colocalization index is equal to 0%) (Fig. S1). *Vice versa*, all mitotic cells are labeled when the exposure time exceeds the length of the G2- and M-phases (the colocalization index is 100%) (Fig. S1). When the population of studied cells is homogeneous in terms of the G2-phase duration, the time lag between 0% and 100% colocalization is equal to the duration of the M-phase. In turn, the time point when half of mitoses are labeled (the colocalization index is 50%) indicates the average duration of G2-phase.

However, we found choanocytes to be heterogeneous in terms of G2-phase length. Few fast-cycling choanocytes managed to pass through the G2-phase during 6 hours of EdU treatment in *L. corallorrhiza* and 9 hours in *H. dujardinii*; at the same time, some mitotic cells were not EdU-positive even after 21 hours of EdU treatment. And while we observed the accumulation of EdU-labeled mitoses by lengthening the EdU exposure, the supposed time lag between 0% and 100% of colocalization was more than 15 hours. Surely, such a time lag cannot be explained by the length of the M-phase. Instead, we expect a relatively broad range of G2-phase lengths to account for the slow accumulation of labeled mitoses. This finding strikingly resembles the case of planarian neoblasts having the median G2-phase of 6 hours despite the presence of a few slowly cycling cells [42].

Nonetheless, we found another option to calculate the length of G2-phase (Table 2). We used cytometry-based assessment of DNA content in single-cell suspensions to distinguish between diploid G0/G1 cells, tetraploid G2/M-cells, and DNA-synthesizing S-phase cells. ACME dissociation and RNAse treatment significantly improved the quality of suspensions compared to the protocol we used before [21], allowing us to perform quantitative analysis of cell cycle distribution curve. Since we showed choanocytes to account for more than 90% of cycling cells, we supposed that the absolute majority of S- and G2/M-cells in the suspensions were choanocytes. By calculating the proportion of S-phase cells to tetraploid G2/M-cells, we found the median G2-phase to last slightly longer than the S-phase (namely, about 15 hours in *H. dujardinii* and 18 hours in *L. corallorrhiza*) (Table 2). Finally, the length of the G1-phase was estimated by subtracting the lengths of S-, G2- and M-phase from the total cell cycle duration; it was nearly 14 hours in *H. dujardinii* and 27 hours in *L. corallorrhiza*. Of course, these estimates are fairly rough; G1- and G2-phases are shown to be the most variable phases of the cell cycle in terms of their duration [43–45]. Since choanocytes are heterogeneous in the length of the G2-phase, they could as well differ in the length of the G1 phase.

### Future perspectives

In this paper, we provide the most current data associated with cell turnover in sponges. This paper serves as the basis for future studies regarding the stem cell system and tissue dynamics of sponges. Here we describe the populations of cycling cells in intact sponges and analyze the cell cycle of the major population of cycling cells, food-entrapping choanocytes. We also introduce aphidicolin as a promising tool for *in vivo* synchronization of cycling cells. Our data complement the existing estimates of cell cycle duration in sponge choanocytes and provide an important insight into the tissue dynamics of sponges. Possibly, the relatively long cell cycle of choanocytes affects not only the tissue renewal but also the dynamics of regeneration. Although the cell cycle kinetics in regenerating sponges is yet to be described, the active transdifferentiation involved in wound healing might be a mechanism to compensate for the long cell cycle duration and achieve fast recovery.

Our study also highlights some important issues to consider when analyzing the cell cycle. The length of the G2-phase, cell heterogeneity, and the growth fraction can drastically affect the estimates and lead to conflicting results. The method introduced here could serve as a reliable and simple technique to comprehensively analyze the cell cycle not only in sponges but also in other animals.

## Materials and methods

### Sampling and laboratory maintenance

Individuals of *Halisarca dujardinii* Johnston, 1842 (Demospongiae, Chondrillida) and *Leucosolenia corallorrhiza* (Haeckel, 1872) (Calcarea, Leucosolenida) were collected in the Kandalaksha Bay in the White Sea near the Pertsov White Sea Biological Station of Lomonosov Moscow State University (66°34′ N, 33°08′ E) in July-August, 2021 and 2022. Sponges were manually collected with the substrate (algae and stones) and kept for two to seven days in laboratory aquariums with running natural seawater of ambient temperature. To avoid excessive inter-individual variability, sponges of the same size and same habitat were used in each experiment. During the experiments, sponges were kept at 8-12°C in plastic Petri dishes with 5 mL of 0.22 μm-filtered seawater (FSW). Sponges were photographed using Nikon Z5 (Nikon, Shinagawa, Japan) digital camera equipped with Zenitar 2,8/60 Macro EA (Zenit) objective.

### Identifying proliferating cell types

DNA-synthesizing cells were revealed by confocal laser scanning microscopy (CLSM) by the incorporation of thymidine analog 5-ethynyl-2’-deoxyuridine (EdU; Lumiprobe, 10540) in sponges vitally stained with cytoplasmic dye CellTracker DeepRed Dye (CellTracker; ThermoFisher Scientific, C34565). CellTracker allowed us to study the morphology of labeled cells and identify cell types by comparison to histological semi-thin sections.

To study the proliferation via CLSM, sponges (three individuals per species) were treated with 100 μM of EdU and 1 mM CellTracker for 6 h (*L. corallorrhiza*) and 12 h (*H. dujardinii*) and fixed with 4% paraformaldehyde in phosphate-buffered saline (4% PFA PBS, Carl Roth 0335.2). Specimens were washed with phosphate-buffered saline and blocking solution (1% BSA, 0.1% gelatin from cold-water fish skin (Sigma-Aldrich G7041), 0.5% Triton X-100 and 0.05% Tween-20 (Sigma-Aldrich P1379) in PBS). EdU was visualized by copper click-chemistry in a solution of 4 mM CuSO_4_, 20 μM Sulfo-Cyanine3 Azide (Lumiprobe A1330) and 20 mg/mL sodium L-ascorbate (Sigma-Aldrich 11140) in PBS. Click-reaction was performed for 2 hours at room temperature. Choanocyte flagella were stained with Mouse Anti-acetylated-α-tubulin primary antibodies (Sigma-Aldrich T6793) diluted 1:1000 and Donkey anti-Mouse IgG Alexa Fluor 488 (ThermoFisher Scientific A21202) secondary antibodies in a concentration of 1 μg/mL. Each round of antibody treatment was performed in blocking solution overnight at +4°C. Nuclei were stained with 4′,6-diamidino-2-phenylindole (DAPI, Acros 202710100) in a concentration of 1 μg/ml for 2 hours at +4°C. Spicules of *L. corallorrhiza* were dissolved with 5% ethylenediaminetetraacetic acid (EDTA) in distilled water for 30 min at room temperature. Finally, specimens were mounted in 90% glycerol (MP Biomedicals 193996) with 2.5% 1,4-diazabicyclo[2.2.2]octane (DABCO, Sigma-Aldrich D27802).

In *H. dujardinii*, we studied the endosome at the periphery of a sponge’s body; in *L. corallorrhiza*, we studied the middle part of an oscular tube, the oscular rim, and diverticula. Three non-overlapping Z-stacks were obtained for each studied region of an individual. Z-stacks were obtained by Nikon A1 (Nikon, Shinagawa, Japan) using excitation lasers of wavelength 405 nm (DAPI), 488 nm (Mouse Anti-acetylated-α-tubulin ABI + DAM IgG Alexa Fluor 488 ABII), 561 nm (EdU + Sulfo-Cyanine3 Azide) and 648 nm (CellTracker). Z-stacks were 20-30 µm thick with a Z-step of 1 µm and contained 1500-4000 cells each.

Z-stack analysis was performed in Fiji (ImageJ) v1.53f51. Cell types were identified by the morphology revealed through CellTracker staining; choanocytes were distinguished by the presence of flagella. Three-dimensional quantification of DNA-synthesizing cells was carried out in Bitplane Imaris v7.2.1. We marked nuclei stained with DAPI or EdU with the “Spots” tool, combining automatic counting via Quality/Intensity Median thresholds with manual correction of resulting segmentation. To count choanocytes separately, we created 3D surfaces judging by the flagella staining around choanocyte chambers/choanoderm and split already existing spots (all labeled cells) between resulting surface objects. For each obtained Z-stack, we estimated the total labeling fraction (as a proportion of all EdU-positive nuclei to all nuclei stained with DAPI), the labeling fraction of choanocytes or mesohyl cells (as a proportion of EdU-positive choanocytes/mesohyl cells to a respective population), and the fraction of EdU-positive choanocytes among all EdU-positive cells.

Since each Z-stack contained numerous cells, we decided to adjust the analysis for the number of registered cells by treating studied parameters as binomially distributed values. Basically, each fraction reflected the probability of a cell being labeled out of “n” trials, where “n” was the number of cells in a Z-stack (about 1500-4000 cells). Thus, we used logistic regression with logit link (binomial regression) to predict population proportions. Because of the nested design of the experiment (each individual was represented by three Z-stacks), we used mixed effects logistic regression with the individual organism as a random effect. Predicted mean values and the corresponding standard errors are reported in the text as the estimates of the studied parameter (labeling fraction, fraction of choanocytes, etc.).

For histological studies, sponges were fixed for 2 hours at 4 °C with 2.5% glutaraldehyde (Electron Microscopy Science, 16020), post-fixed for 1 hour at room temperature with 1% OsO_4_ (Electron Microscopy Science, 19100), dehydrated in the ethanol/acetone series and embedded in Epon/Araldite epoxy embedding media (Electron Microscopy Science, 13940) according to the protocol described earlier [3].

### Estimating the growth fraction

Growth fraction (**GF**) is a fraction of all proliferating cells in a tissue. To estimate GF in sponge tissues, we synchronized cells in the S-phase of the cell cycle and labeled them with EdU. S-phase synchronization was achieved by treating sponges with aphidicolin, a reversible inhibitor of DNA polymerase [22].

The effective concentration of aphidicolin and the expected cell cycle length were estimated during the preliminary experiments. In the final set of experiments, sponges (three individuals per species) were treated with aphidicolin (1 μg/ml in FSW) for 4.5 days (approximately 2× of the expected duration of choanocyte cell cycle). Each day the solution was renewed to avoid the accumulation of metabolites and to sustain stable condition of animals. Sponges were allowed to recover from the S-phase block for 12 h in an aquarium with running natural seawater, as sponges treated with EdU immediately after the incubation showed less proliferating cells than we expected. After the recovery, sponges were treated with 100 μM of EdU for 12 h (*H. dujardinii*) and 6 h (*L. corallorrhiza*) and prepared for CSLM as it was described earlier (see the “Identifying proliferating cell types” section) without using the CellTracker.

Three sponges per species also served as positive controls to ensure that aphidicolin is efficient to inhibit DNA synthesis; these sponges were treated with 100 μM of EdU for 6 hours prior to the end of 4.5 days of aphidicolin treatment (EdU was added directly to the medium containing aphidicolin). Negative controls (sponges treated with 5 μl of DMSO in 5 ml FSW for 4.5 days) were used to set a baseline of the fraction of proliferating cells in presumably suboptimal maintenance conditions.

Three non-overlapping Z-stacks of the endosome (*H. dujardinii*) and the middle part of an oscular tube (*L. corallorrhiza*) were obtained for each aphidicolin-treated individual. One Z-stack was obtained for each control specimen. Stack registration and nuclei counting procedures were described in the “Identifying proliferating cell types” section. The estimates of labeling fractions and the corresponding standard errors were calculated using nested binomial regression with treatment as fixed effect (predictor) and the individual organism as random effect. Wald Chi-square and log-likelihood tests against the model with no fixed effect were performed to report the significance of treatment.

### G2/M-phase analysis

We used Ser10-phosphorylated histone 3 (рН3) monoclonal antibodies to mark late G2- and M-phase cells and analyzed the accumulation of EdU in mitotic cells to estimate the duration of G2-phase. Sponges were treated with 100 μM EdU for 9, 12, 15, 18, and 21 h (*H. dujardinii*) and 6, 9, 12, 15, 18, and 21 h (*L. corallorrhiza*) and prepared for CLSM (see the “Identifying proliferating cells” section) with mitotic cells being labeled by a combination of Rabbit Anti-phospho-histone H3, Sigma-Aldrich H0412 primary antibodies (diluted 1:1000) and Rabbit IgG Alexa Fluor 647, ThermoFisher Scientific A-31573 secondary antibodies (in a concentration of 1 μg/mL). Three individuals of *H. dujardinii* were used per time point; in the case of *L. corallorrhiza*, we used a total of three large multioscular sponges, cutting off a single oscular tube at any given time point. Three non-overlapping Z-stacks of the endosome (*H. dujardinii*) and the middle part of an oscular tube (*L. corallorrhiza*) were obtained for each individual of the time point.

Subsequent Z-stack analysis was focused on choanocytes since they comprised the absolute majority of pH3-labeled cells. For each individual, we analyzed the orientation of mitotic plates, the mitotic fraction (the proportion of pH3-positive choanocytes to all DAPI-stained choanocyte nuclei), and the colocalization index (the proportion of EdU-labeled pH3-positive choanocytes to all pH3-positive choanocytes). We assumed that choanocytes were homogeneous in terms of the duration of the G2-phase; in such case, the time point with 50% mitoses being labeled should correspond to the average length of the G2-phase [23,42]. In turn, the period between time points of 0% and 100% colocalization index should correspond to the average length of the M-phase (Fig. S1).

We analyzed the effect of the incubation on the fraction of labeled mitoses by performing nested binomial regression with treatment time as the fixed effect (numeric predictor) and the individual organism as the random effect. Wald Chi-square and log-likelihood tests against the model with only the random effect were performed to report the significance of the fixed effect. Predicted values and the corresponding standard errors are provided in the text as the estimates for the fraction of labeled mitoses at each time point of the experiment. Since the binomial distribution results in non-linear effect of the predictor (treatment time) on the response value (the fraction of labeled choanocytes), the average increase in the fraction of labeled mitoses was retrieved by computing the average marginal effect.

### Cell cycle dynamics

The estimation of cell cycle dynamics (namely, the duration of the cell cycle ***T_C_***) was performed according to Sanders’ mathematical model [34,46]. This model implies the analysis of EdU incorporation during a short period of time. Basically, we expect more EdU-positive cells to appear during a continuous EdU treatment: some cells enter the S-phase and become labeled, whereas others leave the S-phase and retain the label. If so, the fraction of labeled cells (***F***) is determined by the treatment time (***T_E_***) and the unknown lengths of the S-phase (***T_S_***) and the cell cycle (***T_C_***) (equation 1):

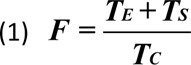

Given that two different treatment times (***T_E1_*** and ***T_E2_***) are used, we can calculate the unknown lengths (equation 2):

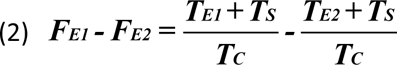

Solving the equation (2) for ***T_C_*** results in a formula (3):

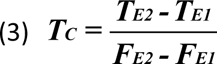

However, since not all cells are equally proliferative (there could possibly be quiescent or non-proliferative cells), we need to take into account the growth fraction (**GF**), the fraction of all cycling cells in a given tissue. GF was assumed to be equal to the choanocyte labeling fraction after aphidicolin-mediated S-phase synchronization. The final equation for calculating the cell cycle length is (4):

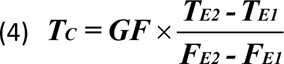

To avoid the accumulation of EdU-positive cells due to the division of labeled cells, the maximum treatment time should be less than the duration of the G2-phase. Considering the results of the G2-phase duration analysis (see the “G2/M-phase analysis” section), we treated sponges with 100 μM EdU for ***T_E1_***=2 h and ***T_E2_***=6 h. Three individuals of *H. dujardinii* were used per time point; in the case of *L. corallorrhiza*, we used a total of three large multioscular sponges, cutting off a single oscular tube at given time points. Sponge tissues were fixed and prepared for CLSM as was described earlier (see the “Identifying proliferating cell types” section) without using the CellTracker. We studied the endosome of *H. dujardinii* and the middle part of the oscular tube of *L. corallorrhiza*, recording three non-overlapping Z-stacks for each individual.

We calculated the labeling fraction of choanocytes at each time point with the corresponding standard errors by performing nested binomial regression with treatment time as the fixed effect (numeric predictor) and the individual organism as the random effect. Wald Chi-square and log-likelihood tests against the model with no fixed effect were performed to report the significance of treatment time. Since the binomial distribution results in non-linear effect of the predictor (treatment time) on the response value (the fraction of labeled choanocytes), the fraction of choanocytes becoming labeled each hour was retrieved by computing the average marginal effect. Finally, the duration of the cell cycle (***T_C_***) was calculated according to the equation (4) using standard rules for calculating the propagation of uncertainty. Thus, the estimates for the duration of the cell cycle (***T_C_***) are presented in the text as mean ± standard error.

The lengths of S- and M-phases (***T_S_*** and ***T_M_***, respectively) were calculated according to the assumption that the percentage of cells in each phase of the cell cycle corresponds to the proportion of each phase to the total duration of the cell cycle (5 and 6):

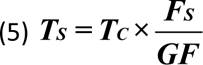

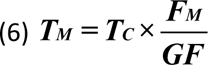

The fraction of choanocytes in the S-phase at any given time point ***F_S_*** was estimated as the fraction of labeled choanocytes at T=0 h (i.e. the intercept of the fitted binomial regression model). The fraction of mitotic choanocytes ***F_M_*** was calculated in the G2/M-phase analysis experiments as the mean fraction of pH3-labeled choanocytes among all studied time points.

### Flow cytometry: cell cycle distribution

Another option to analyze the cell cycle (rather than using certain specific markers) is provided by the quantitative DNA staining utilized in flow cytometry. Proper staining with DNA intercalators such as propidium iodide (PI) reveals the DNA content in each cell of a suspension and helps to differ between tetraploid (G2/M) and diploid (G1/G0) cells with S-phase cells showing an intermediate amount of DNA [47].

Cell suspensions were obtained by adapting the ACME dissociation for sponge tissues [48]. Sponges (three individuals per species) were fixed in 10 ml of ACME solution (distilled H2O, methanol, glacial acetic acid and glycerol in a proportion of 13:3:2:2), homogenized for 10 min with a tissue homogenizer RT-2 (USSR) and stored at +4°C for 2 months. Suspensions were washed with PBS and blocking solution (1% BSA (MP Biomedicals 0216006980), 0.1% gelatin from cold-water fish skin (Sigma-Aldrich G7041), 0.5% Triton X-100 and 0.05% Tween-20 (Sigma-Aldrich P1379) in PBS) several times and stained with propidium iodide (P1304MP, ThermoFisher Scientific) in a concentration of 2 µg/ml. Staining was performed overnight at +4°C with an addition of 10 µg/ml RNAse A from bovine pancreas (Sigma-Aldritch, R6513) to eliminate possible fluorescence of PI-stained RNA.

FACSAria SORP cell sorter (BD Biosciences, USA) with an excitation laser wavelength of 561 nm was used to record 50 000 events per suspension. The data was processed in FlowJo V10 (BD Biosciences). Primary gates to differ between debris and intact cells were set according to forward (FCS-A) versus side scatter (SSC-A) parameters; secondary gating to distinguish single cells was performed according to FSC-W versus FSC-H parameters (Fig. S2). The final cell cycle curves were visualized in a form of a single-parameter histogram based on the intensity of PI fluorescence.

Cell cycle distribution was assessed by FlowJo CellCycle plugin approximating the DNA content curve to a sum of two Gaussian distributions – namely, G0/G1 and G2/M cells; events between G0/G1 and G2/M cells were regarded as S-phase cells. We used standard parameters of the Watson model, constraining the position of the G2/M peak according to the position of the G0/G1 cells. No CV constraint was performed. Fractions of G1/G0, G2/M, and S-cells are presented in the text as mean values ± standard deviation.

We used the mean proportion of G2/M-cells to S-cells and the calculated the length of S-phase (see the “Cell cycle dynamics” section) to predict the average total length of the G2-phase: 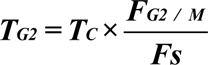 and qualitatively estimate the length of G1 phase: ***T_G1_ = T_C_ - T_S_ - T_G2_ - T_M_***.

### Statistical analysis and plotting

Statistical analysis was conducted in Rstudio using lme4 package to perform nested logistic regression and emmeans package to return predictions, the corresponding standard errors and 95% confidence intervals. Wald Chi-square tests of the computed models were conducted using type III Anova function provided in car package. Log-likelihood tests were conducted using anova test against the models with no fixed effects. Average marginal effects were computed using margins library. Plotting was performed using ggplot2 library. R scripts are provided in Supporting Information.

## Author Contributions

AL and NM designed the study. NM and AL collected the material and prepared the specimens for CLSM. NM collected and prepared the specimens for flow cytometry. Quantitative and statistical analysis of CLSM and flow cytometry data was performed by NM. AL and NM prepared the text and the figures. All authors reviewed and approved the final manuscript.

## Abbreviations

CLSM: confocal laser scanning microscopy
CellTracker: CellTrackerTM DeepRed Dye
DAPI: 4′,6-diamidino-2-phenylindole
DMSO: dimethyl sulfoxide, EdU, 5-ethynyl-2’-deoxyuridine
FG1: the fraction of cells in the G1 phase
FG2: the fraction of cells in the G2 phase
FM: the fraction of cells in the M phase
FS: the fraction of cells in the S phase
FSW: filtered seawater
GF: growth fraction
PBS: phosphate-buffered saline
PFA: paraformaldehyde
pH3: Ser10-phosphorylated histone 3
PI: propidium iodide
TC: the total cell cycle duration
TG1: the length of the G1 phase
TG2: the length of the G2 phase
TM: the length of the M phase
TS: the length of the S phase

## Acknowledgements

The authors acknowledge the support of Lomonosov Moscow State University Program of Development (FACSAria SORP flow cytometer/sorter, Nikon A1 CLSM) and Center of microscopy WSBS MSU. Authors sincerely thank Darya Potashnikova (Lomonosov Moscow State University) for operating the flow cytometer, Kseniia Skorentseva for proofreading the manuscript, Fyodor Bolshakov for providing us with digital photos of intact sponges, Elena Voronezhskaya (Koltsov Institute of Developmental Biology) and Igor Kosevich (Lomonosov Moscow State University) for helpful tips and advice.

## Data availability

Confocal Z-stacks and raw flow cytometry data supporting the conclusions of this article are available in the Mendeley Data repository under the following DOI: 10.17632/7fbv46wsnv.1 (*H. dujardinii*) and 10.17632/r3ydrmryzg.1 (*L. corallorrhiza*). The results of the quantitative analysis of proliferation are presented as .xlsx table (raw data) and R script (statistical analysis procedures) in the Supplementary information.

