## Supplementary material for "An experimental approach in analyzing the cell cycle dynamics of food-entrapping cells of sponges": cell cycle statistics.R

```
####importing libraries
library(readxl)
library(ggplot2)
theme_set(theme_bw())
library(scales)
library(lme4)
library(car)
library(lsmeans)
library(margins)
library(pals)

inverse_logit = function(x){
  exp(x)/(1+exp(x))
}


#importing data
cell_cycle_data <- read_excel("D:/MNP/cell cycle data.xlsx")
cell_cycle_data <- read_excel("G:/?????? ??????????/?????????/MNP/cell cycle data.xlsx")
df <- as.data.frame(cell_cycle_data)


##########regular proliferation

proliferation_data <- read_excel("G:/?????? ??????????/?????????/MNP/intact proliferation.xlsx")
df2 <- as.data.frame(proliferation_data)

###hd
dfhd <- subset(df2, df2$sp=="Hd")

dfhd$dapimes <- dfhd$dapi-dfhd$dapich
dfhd$edumes <- dfhd$edu-dfhd$educh

#total labeled cells
hd_fit <-  glmer(cbind(edu, dapi-edu)~1+(1|ind), data=dfhd, family='binomial')
data.frame(emmeans(hd_fit, '1', type='response'))*100


#labeled choanocytes
hd_fit_ch <-  glmer(cbind(educh, dapich-educh)~1+(1|ind), data=dfhd, family='binomial')
data.frame(emmeans(hd_fit_ch, '1', type='response'))*100

#labeled mesohyl
hd_fit_mes <-  glmer(cbind(edumes, dapimes-edumes)~1+(1|ind), data=dfhd, family='binomial')
data.frame(emmeans(hd_fit_mes, '1', type='response'))*100


###lc
dflc <- subset(df2, df2$sp=="Lc")

dflc$dapimes <- dflc$dapi-dflc$dapich
dflc$edumes <- dflc$edu-dflc$educh

#total labeled cells
lc_fit <-  glmer(cbind(edu, dapi-edu)~1+(1|ind), data=dflc, family='binomial')
data.frame(emmeans(lc_fit, '1', type='response'))*100


#labeled choanocytes
lc_fit_ch <-  glmer(cbind(educh, dapich-educh)~1+(1|ind), data=dflc, family='binomial')
data.frame(emmeans(lc_fit_ch, '1', type='response'))*100

#labeled mesohyl
lc_fit_mes <-  glmer(cbind(edumes, dapimes-edumes)~1+(1|ind), data=dflc, family='binomial')
data.frame(emmeans(lc_fit_mes, '1', type='response'))*100


############# Growth fraction
#####Lc
dfpflc <- subset(df, df$exp=="pf"&df$sp=="Lc")
dfpflc$ind=as.factor(c(1,1,1,2,2,2,3,3,3,4,5,6))
dfpflc$t = as.factor(dfpflc$t)

####performing the GLMM for the total labeling fraction
pf_lc_fit <-  glmer(cbind(edu, dapi-edu)~t+(1|ind), data=dfpflc, family='binomial')
summary(pf_lc_fit)

#checking the effects
#fitting reduced model
pf_lc_no_fit <- glmer(cbind(edu, dapi-edu)~1+(1|ind), data=dfpflc, family='binomial')
summary(pf_lc_no_fit)

anova(pf_lc_fit, pf_lc_no_fit) #loglik test: treatment is significant

#returning estimates and SE
lcpftotal <- data.frame(emmeans(pf_lc_fit, 't', type='response'))

####performing the GLMM for choanocyte labeling fraction
pf_lc_fit_ch <-  glmer(cbind(educh, dapich-educh)~t+(1|ind), data=dfpflc, family='binomial')
summary(pf_lc_fit_ch)

#reduced model
pf_lc_no_fit_ch <- glmer(cbind(educh, dapich-educh)~1+(1|ind), data=dfpflc, family='binomial')
anova(pf_lc_fit_ch, pf_lc_no_fit_ch) #loglik test: treatment is significant

#returning estimates and SE
lcpf <- data.frame(emmeans(pf_lc_fit_ch, 't', type='response'))

####performing the GLMM for mesohyl labeling fraction
pf_lc_fit_mes <-  glmer(cbind(edumes, dapimes-edumes)~t+(1|ind), data=dfpflc, family='binomial')
summary(pf_lc_fit_mes)

#reduced model
pf_lc_no_fit_mes <-  glmer(cbind(edumes, dapimes-edumes)~1+(1|ind), data=dfpflc, family='binomial')
anova(pf_lc_fit_mes, pf_lc_no_fit_mes) #loglik test: treatment is significant

#returning estimates and SE
lcpfmes <- data.frame(emmeans(pf_lc_fit_mes, 't', type='response'))

####performing the GLMM for the fraction of choanocytes among all edu-positive cells
pf_lc_ch.vs.mes <-  glmer(cbind(educh, edu-educh)~t+(1|ind), data=dfpflc, family='binomial')
summary(pf_lc_ch.vs.mes)

#checking the effects
pf_lc_ch.vs.mes_red <-  glmer(cbind(educh, edu-educh)~1+(1|ind), data=dfpflc, family='binomial')
anova(pf_lc_ch.vs.mes, pf_lc_ch.vs.mes_red )#logLik: treatment is NOT significant

#returning estimates and SE
emmeans(pf_lc_ch.vs.mes , 't', type='response')


#######plotting
ggplot()+
  geom_boxplot(data=lcpf, aes(x=t, middle=prob, lower=asymp.LCL, upper=asymp.UCL, ymin=asymp.LCL, ymax=asymp.UCL), stat='identity', width=0.15, lwd=1, position=position_nudge(0.2))+
  geom_point(data=dfpflc, aes(x=t, y=educh/dapich, col=ind), stroke=0, size=4, position=position_jitternudge(x=0.2, y=0, width=0.03, nudge.from = 'jittered'), alpha=0.5)+
  
  geom_boxplot(data=lcpftotal, aes(x=t, middle=prob, lower=asymp.LCL, upper=asymp.UCL, ymin=asymp.LCL, ymax=asymp.UCL), stat='identity', width=0.15, lwd=1)+
  geom_point(data=dfpflc, aes(x=t, y=edu/dapi, col=ind), stroke=0, size=4, position=position_jitternudge(x=0, y=0, width=0.03, nudge.from = 'jittered'), alpha=0.5)+
  
  geom_boxplot(data=lcpfmes, aes(x=t, middle=prob, lower=asymp.LCL, upper=asymp.UCL, ymin=asymp.LCL, ymax=asymp.UCL), stat='identity', width=0.15, lwd=1, position=position_nudge(-0.2))+
  geom_point(data=dfpflc, aes(x=t, y=edumes/dapimes, col=ind), stroke=0, size=4, position=position_jitternudge(x=-0.2, y=0, width=0.03, nudge.from = 'jittered'), alpha=0.5)+
  
  ylim(0,0.7)+
  theme(legend.position="none")


#####hd
dfpfhd <- subset(df, df$exp=="pf"&df$sp=="Hd")
dfpfhd$ind=as.factor(c(1,1,1,2,2,2,3,3,3,3,4,4,5,6))
dfpfhd$t = as.factor(dfpfhd$t)

####performing the GLMM for the total labeling fraction
pf_hd_fit <-  glmer(cbind(edu, dapi-edu)~t+(1|ind), data=dfpfhd, family='binomial')
summary(pf_hd_fit)

#reduced model
pf_hd_no_fit <-  glmer(cbind(edu, dapi-edu)~1+(1|ind), data=dfpfhd, family='binomial')
anova(pf_hd_fit, pf_hd_no_fit) #logLik: treatment is significant

#returning estimates and SE
hdpftotal <- data.frame(emmeans(pf_hd_fit, 't', type='response'))

####performing the GLMM for choanocyte labeling fraction
pf_hd_fit_ch <-  glmer(cbind(educh, dapich-educh)~t+(1|ind), data=dfpfhd, family='binomial')
summary(pf_hd_fit_ch)

#reduced model
pf_hd_no_fit_ch <-  glmer(cbind(educh, dapich-educh)~1+(1|ind), data=dfpfhd, family='binomial')
anova(pf_hd_fit_ch, pf_hd_no_fit_ch) #logLik: treatment is significant

#returning estimates and SE
hdpf <- data.frame(emmeans(pf_hd_fit_ch, 't', type='response'))

####performing the GLMM for mesohyl labeling fraction
pf_hd_fit_mes <-  glmer(cbind(edumes, dapimes-edumes)~t+(1|ind), data=dfpfhd, family='binomial')
summary(pf_hd_fit_mes)

#reduced model
pf_hd_no_fit_mes <-  glmer(cbind(edumes, dapimes-edumes)~1+(1|ind), data=dfpfhd, family='binomial')
anova(pf_hd_fit_mes, pf_hd_no_fit_mes) #logLik: treatment is significant

#returning estimates and SE
hdpfmes <- data.frame(emmeans(pf_hd_fit_mes, 't', type='response'))

####performing the GLMM for the fraction of choanocytes among all edu-positive cells
pf_hd_ch.vs.mes <-  glmer(cbind(educh, edu-educh)~t+(1|ind), data=dfpfhd, family='binomial')
summary(pf_hd_ch.vs.mes)

#checking the effects
pf_hd_ch.vs.mes_red <-  glmer(cbind(educh, edu-educh)~1+(1|ind), data=dfpfhd, family='binomial')
anova(pf_hd_ch.vs.mes, pf_hd_ch.vs.mes_red) #logLik: treatment is NOT significant

#returning estimates and SE
emmeans(pf_hd_ch.vs.mes, 't', type='response')

ggplot()+
  geom_boxplot(data=hdpf, aes(x=t, middle=prob, lower=asymp.LCL, upper=asymp.UCL, ymin=asymp.LCL, ymax=asymp.UCL), stat='identity', width=0.15, lwd=1, position=position_nudge(0.2))+
  geom_point(data=dfpfhd, aes(x=t, y=educh/dapich, col=ind), stroke=0, size=4, position=position_jitternudge(x=0.2, y=0, width=0.03, nudge.from = 'jittered'), alpha=0.5)+
  
  geom_boxplot(data=hdpftotal, aes(x=t, middle=prob, lower=asymp.LCL, upper=asymp.UCL, ymin=asymp.LCL, ymax=asymp.UCL), stat='identity', width=0.15, lwd=1)+
  geom_point(data=dfpfhd, aes(x=t, y=edu/dapi, col=ind), stroke=0, size=4, position=position_jitternudge(x=0, y=0, width=0.03, nudge.from = 'jittered'), alpha=0.5)+
  
  geom_boxplot(data=hdpfmes, aes(x=t, middle=prob, lower=asymp.LCL, upper=asymp.UCL, ymin=asymp.LCL, ymax=asymp.UCL), stat='identity', width=0.15, lwd=1, position=position_nudge(-0.2))+
  geom_point(data=dfpfhd, aes(x=t, y=edumes/dapimes, col=ind), stroke=0, size=4, position=position_jitternudge(x=-0.2, y=0, width=0.03, nudge.from = 'jittered'), alpha=0.5)+
  
  ylim(0,0.7)+
  theme(legend.position="none")


###########S PHASE ANALYSIS
#####Lc
dflcacc <- subset(df, df$sp=="Lc"&df$exp=="acc")
dflcacc$t <- as.numeric(dflcacc$t)
dflcacc$ind <- as.factor(rep(c(1,2,3,1,2,3), each=3))

#choanocyte labeling fraction
acc_lc_fit <- glmer(cbind(educh, dapich-educh)~t+(1|ind), data=dflcacc, family='binomial')
summary(acc_lc_fit)

#reduced model
acc_lc_fit_red <- glmer(cbind(educh, dapich-educh)~1+(1|ind), data=dflcacc, family='binomial')
anova(acc_lc_fit,acc_lc_fit_red) #logLik: time is significant


#returning estimates and SE
lcacc <- data.frame(emmeans(acc_lc_fit, at=list(t=c(0:7)), 't',type='response'))


summary(margins(acc_lc_fit, type='response', data=dflcacc))


#############plotting
ggplot()+
  geom_line(data=lcacc, aes(x=t, y=prob), lwd=1.5, alpha=0.7)+
  geom_ribbon(data=lcacc, aes(x=t, ymax=asymp.UCL, ymin=asymp.LCL), alpha=0.05)+
  geom_point(data=dflcacc, aes(x=t, y=educh/dapich, col=ind), size=8, alpha=0.7, stroke=0, position=position_jitter(0.2))+
  scale_x_continuous(breaks=c(0,2,4,6), limits=c(0,7))+
  ylim(0.05, 0.25)+
  theme(legend.position="none")


#####Hd
dfhdacc <- subset(df, df$sp=="Hd"&df$exp=="acc")
dfhdacc$t <- as.numeric(dfhdacc$t)
dfhdacc$ind <- as.factor(rep(c(1,2,3,4,5,6), each=3))
dfhdacc$stack <- as.factor(c(1:18))

#choanocyte labeling fraction
acc_hd_fit <- glmer(cbind(educh, dapich-educh)~t+(1|ind), data=dfhdacc, family='binomial')
summary(acc_hd_fit)

acc_hd_fit_red <- glmer(cbind(educh, dapich-educh)~1+(1|ind), data=dfhdacc, family='binomial')
anova(acc_hd_fit_red,acc_hd_fit) #logLik: time is significant

#returning estimates and SE
hdacc <- data.frame(emmeans(acc_hd_fit, at=list(t=c(0:7)), 't', type='response'))

summary(margins(acc_hd_fit, type='response', data=dfhdacc))


#############plotting
ggplot()+
  geom_line(data=hdacc, aes(x=t, y=prob), lwd=1.5, alpha=0.7)+
  geom_ribbon(data=hdacc, aes(x=t, ymax=asymp.UCL, ymin=asymp.LCL), alpha=0.05)+
  geom_point(data=dfhdacc, aes(x=t, y=educh/dapich, col=ind), size=8, alpha=0.7, stroke=0, position=position_jitter(0.2))+
  scale_x_continuous(breaks=c(0,2,4,6), limits=c(0,7))+
  ylim(0.05, 0.25)+
  theme(legend.position="none")


#######G2M phase analysis
#####Lc
dflcmit <- subset(df, df$exp=="coloc"&df$sp=="Lc")
dflcmit$ind <- as.factor(dflcmit$ind)
dflcmit$ind2 <- as.factor(c(rep(c(1:16), each=3)))
dflcmit$t=as.numeric(dflcmit$t)

####mitotic fraction
mit_lc_fit <- glmer(cbind(ph3, dapi-ph3)~1+(1|ind), data=dflcmit, family='binomial')
summary(mit_lc_fit)

lcmit <- data.frame(emmeans(mit_lc_fit, '1',type='response'))
lcmit$prob[1]*100
lcmit$SE[1]*100

####mitotic fraction in choanoderm
mitch_lc_fit <- glmer(cbind(ph3ch, dapich-ph3ch)~1+(1|ind), data=dflcmit, family='binomial')
summary(mitch_lc_fit)

lcmitch <- data.frame(emmeans(mitch_lc_fit, '1',type='response'))
lcmitch$prob[1]*100
lcmitch$SE[1]*100

####fraction of choanocytes among ph3 cells
mitfract_lc_fit <- glmer(cbind(ph3ch, ph3-ph3ch)~1+(1|ind), data=dflcmit, family='binomial')
summary(mitfract_lc_fit)

lcmitfract <- data.frame(emmeans(mitfract_lc_fit, '1',type='response'))
lcmitfract$prob[1]*100
lcmitfract$SE[1]*100


#####hd
dfhdmit <- subset(df, df$exp=="coloc"&df$sp=="Hd")
dfhdmit$ind <- c(rep(c(1,2,3,4,5,6,7,8,9,10,11), each=3), 11,12, rep(c(13,14,15), each=3))
dfhdmit$ind <- as.factor(dfhdmit$ind)
dfhdmit$t=as.numeric(dfhdmit$t)

####mitotic fraction
mit_hd_fit <- glmer(cbind(ph3, dapi-ph3)~1+(1|ind), data=dfhdmit, family='binomial')
summary(mit_hd_fit)

hdmit <- data.frame(emmeans(mit_hd_fit, '1',type='response'))
hdmit$prob[1]*100
hdmit$SE[1]*100

####mitotic fraction in choanoderm
mitch_hd_fit <- glmer(cbind(ph3ch, dapich-ph3ch)~1+(1|ind), data=dfhdmit, family='binomial')
summary(mitch_hd_fit)

hdmitch <- data.frame(emmeans(mitch_hd_fit, '1',type='response'))
hdmitch$prob[1]*100
hdmitch$SE[1]*100

####fraction of choanocytes among ph3 cells
mitfract_hd_fit <- glmer(cbind(ph3ch, ph3-ph3ch)~1+(1|ind), data=dfhdmit, family='binomial')
summary(mitfract_hd_fit)

hdmitfract <- data.frame(emmeans(mitfract_hd_fit, '1',type='response'))
hdmitfract$prob[1]*100
hdmitfract$SE[1]*100

####colocalization analysis
####Lc
lc_coloc_fit <- glmer(cbind(colocch, ph3ch-colocch)~t+(1|ind), data=dflcmit, family='binomial', nAGQ=100)
summary(lc_coloc_fit)

#loglikelihood test
lc_coloc_fit_red <- glmer(cbind(colocch, ph3ch-colocch)~1+(1|ind), data=dflcmit, family='binomial')
summary(lc_coloc_fit_red)

anova(lc_coloc_fit, lc_coloc_fit_red) #time is significant

lc_coloc <- data.frame(emmeans(lc_coloc_fit, at=list(t=c(0:24)), 't',type='response'))
lc_coloc*100

summary(margins(lc_coloc_fit, type='response', data=dflcmit))


#############plotting
ggplot()+
  geom_line(data=lc_coloc, aes(x=t, y=prob), lwd=1.5, alpha=0.7)+
  geom_ribbon(data=lc_coloc, aes(x=t, ymax=asymp.UCL, ymin=asymp.LCL), alpha=0.1)+
  geom_point(data=dflcmit, aes(x=t, y=colocch/ph3ch, col=ind, size=ph3ch), alpha=0.7, stroke=0, position=position_jitter(0.2))+
  scale_x_continuous(breaks=c(3,6,9,12,15,18,21,24), limits=c(3,24))+
  scale_size(range=c(4,10))+
  scale_colour_manual(values=as.vector(glasbey(3)))

###Hd
hd_coloc_fit <- glmer(cbind(colocch, ph3ch-colocch)~t+(1|ind), data=dfhdmit, family='binomial')
summary(hd_coloc_fit)

#loglikelihood test
hd_coloc_fit_red <- glmer(cbind(colocch, ph3ch-colocch)~1+(1|ind), data=dfhdmit, family='binomial')
summary(hd_coloc_fit_red)

anova(hd_coloc_fit, hd_coloc_fit_red) #time is significant

hd_coloc <- data.frame(emmeans(hd_coloc_fit, 't', at=list(t=c(5:24)), type='response'))
hd_coloc*100

summary(margins(hd_coloc_fit, type='response', data=dfhdmit))

#############plotting
ggplot()+
  geom_line(data=hd_coloc, aes(x=t, y=prob), lwd=1.5, alpha=0.7)+
  geom_ribbon(data=hd_coloc, aes(x=t, ymax=asymp.UCL, ymin=asymp.LCL), alpha=0.05)+
  geom_point(data=dfhdmit, aes(x=t, y=colocch/ph3ch, col=ind, size=ph3ch), alpha=0.7, stroke=0, position=position_jitter(0.2))+
  scale_x_continuous(breaks=c(6,9,12,15,18,21,24), limits=c(6,24))+
  scale_colour_manual(values=as.vector(glasbey(16)))+
  scale_size(range=c(4,10))


#########CELL CYCLE CALCULATION
###########Hd

#length of the cell cycle
hd.Tc <- hdpf$prob[1]*(6-2)/(hdacc$prob[7]-hdacc$prob[3])

#difference between T6 and T2
hd.dF <- hdacc$prob[7]-hdacc$prob[3]

#SE for difference between T6 and T2
hd.F6.error <-  hdacc$SE[7]
hd.F2.error <- hdacc$SE[3]
hd.dF.error <- sqrt(hd.F6.error^2+hd.F2.error^2)

#SE for growth fraction
hd.pf <- hdpf$prob[1]
hd.pf.error <- hdpf$SE[1]

#relative error of Tc
hd.Tc.rel.error <- sqrt((hd.pf.error/hd.pf)^2+(hd.dF.error/hd.dF)^2)

#SE for hd.Tc
hd.Tc.error <- hd.Tc.rel.error*hd.Tc

#Tc
hd.Ts <- hd.Tc*hdacc$prob[1]/hd.pf

hd.Fs.adj <- hdacc$prob[1]/hd.pf
hd.Fs.adj.error <- hd.Fs.adj*sqrt((hdacc$SE[1]/hdacc$prob[1])^2+(hd.pf.error/hd.pf)^2)
hd.Ts.error <- hd.Ts*sqrt((hd.Fs.adj.error/hd.Fs.adj)^2+(hd.Tc.error/hd.Tc)^2)

#Tm
hd.Tm <- hd.Tc*hdmit$prob[1]/hd.pf

hd.Fm.adj <- hdmit$prob[1]/hd.pf
hd.Fm.adj.error <- hd.Fm.adj*sqrt((hdmit$SE[1]/hdmit$prob[1])^2+(hd.pf.error/hd.pf)^2)
hd.Tm.error <- hd.Tm*sqrt((hd.Fm.adj.error/hd.Fm.adj)^2+(hd.Tc.error/hd.Tc)^2)


###########lc

#length of the cell cycle
lc.Tc <- lcpf$prob[1]*(6-2)/(lcacc$prob[7]-lcacc$prob[3])

#difference between T6 and T2
lc.dF <- lcacc$prob[7]-lcacc$prob[3]

#SE for difference between T6 and T2
lc.F6.error <-  lcacc$SE[7]
lc.F2.error <- lcacc$SE[3]
lc.dF.error <- sqrt(lc.F6.error^2+lc.F2.error^2)

#SE for growth fraction
lc.pf <- lcpf$prob[1]
lc.pf.error <- lcpf$SE[1]

#relative error of Tc
lc.Tc.rel.error <- sqrt((lc.pf.error/lc.pf)^2+(lc.dF.error/lc.dF)^2)

#SE for lc.Tc
lc.Tc.error <- lc.Tc.rel.error*lc.Tc

#Tc
lc.Ts <- lc.Tc*lcacc$prob[1]/lc.pf

lc.Fs.adj <- lcacc$prob[1]/lc.pf
lc.Fs.adj.error <- lc.Fs.adj*sqrt((lcacc$SE[1]/lcacc$prob[1])^2+(lc.pf.error/lc.pf)^2)
lc.Ts.error <- lc.Ts*sqrt((lc.Fs.adj.error/lc.Fs.adj)^2+(lc.Tc.error/lc.Tc)^2)

#Tm
lc.Tm <- lc.Tc*lcmit$prob[1]/lc.pf

lc.Fm.adj <- lcmit$prob[1]/lc.pf
lc.Fm.adj.error <- lc.Fm.adj*sqrt((lcmit$SE[1]/lcmit$prob[1])^2+(lc.pf.error/lc.pf)^2)
lc.Tm.error <- lc.Tm*sqrt((lc.Fm.adj.error/lc.Fm.adj)^2+(lc.Tc.error/lc.Tc)^2)
```
