## Supplementary figures and images for "An experimental approach in analyzing the cell cycle dynamics of food-entrapping cells of sponges"

### Figure S1.png

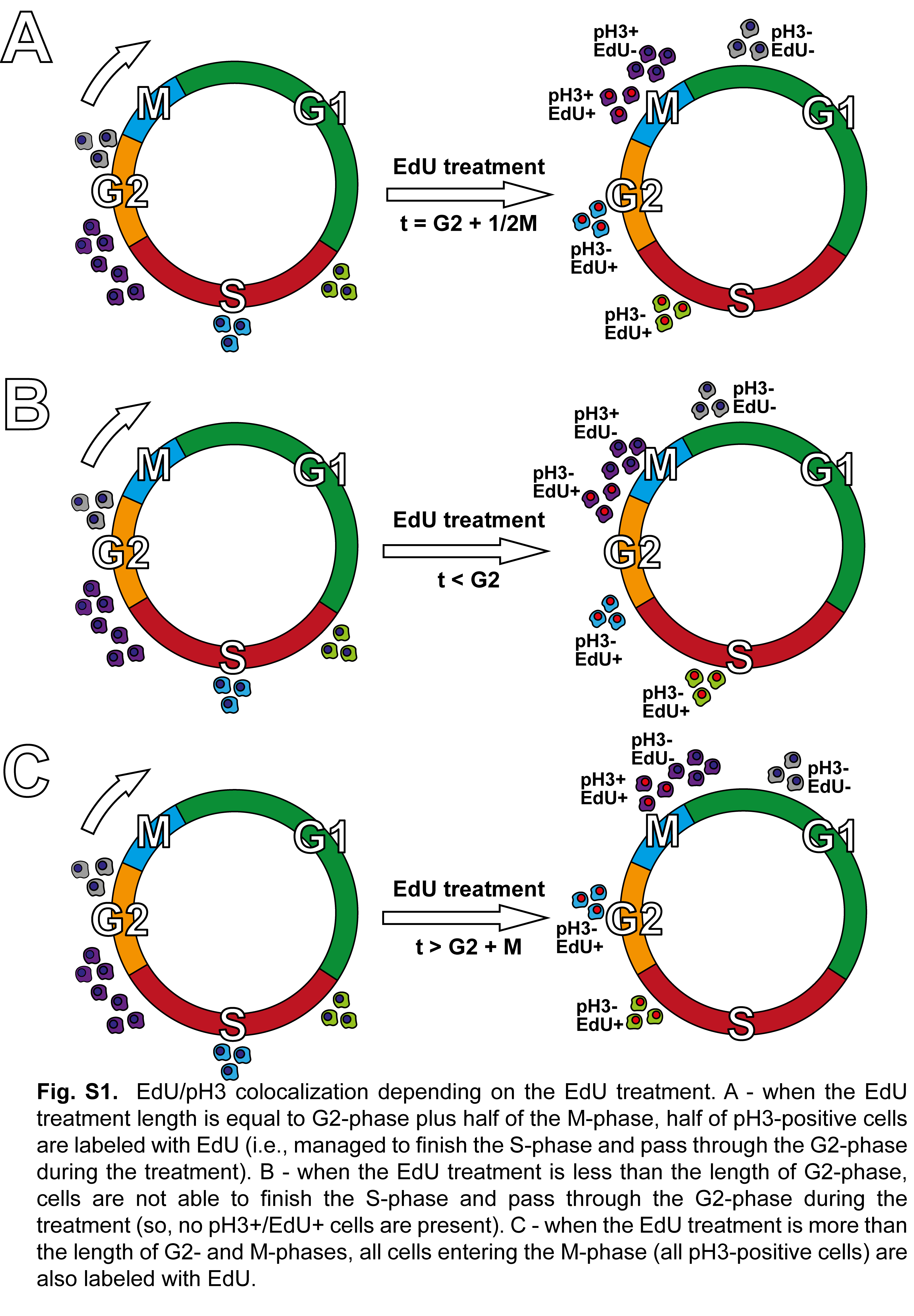

### Figure S2.png

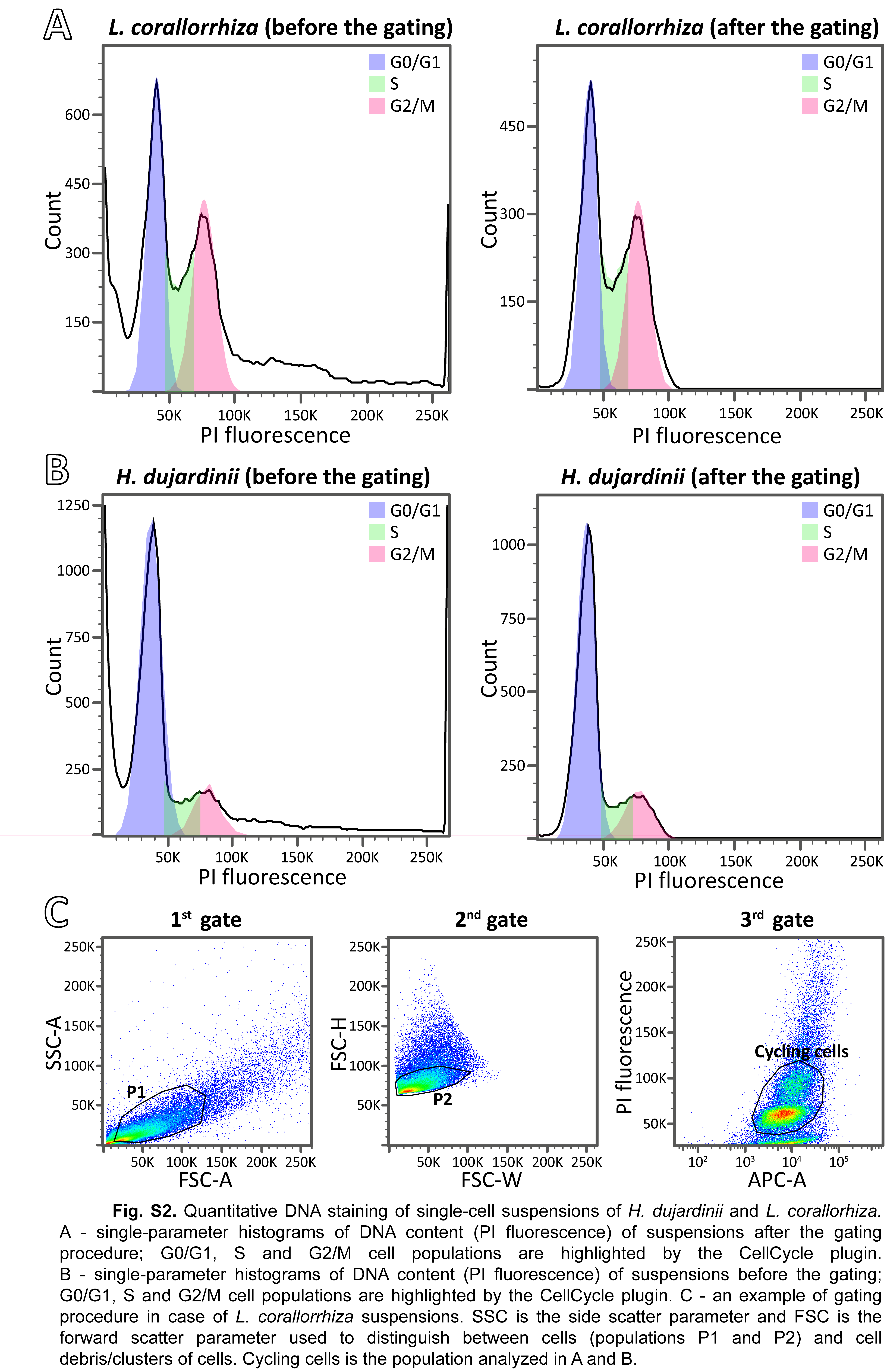
